# LYCEUM: Learning to call copy number variants on low coverage ancient genomes

**DOI:** 10.1101/2024.10.28.620589

**Authors:** Mehmet Alper Yılmaz, Ahmet Arda Ceylan, Gün Kaynar, A. Ercüment Çiçek

## Abstract

Copy number variants (CNVs) are pivotal in driving phenotypic variation that facilitates species adaptation. They are significant contributors to various disorders, making ancient genomes crucial for uncovering the genetic origins of disease susceptibility across populations. However, detecting CNVs in ancient DNA (aDNA) samples poses substantial challenges due to several factors: (i) aDNA is often highly degraded; (ii) contamination from microbial DNA and DNA from closely related species introduce additional noise into sequencing data; and finally, (iii) the typically low coverage of aDNA renders accurate CNV detection particularly difficult. Conventional CNV calling algorithms, which are optimized for high coverage read-depth signals, underperform under such conditions. To address these limitations, we introduce LYCEUM, the first machine learning-based CNV caller for aDNA. To overcome challenges related to data quality and scarcity, we employ a two-step training strategy. First, the model is pre-trained on whole genome sequencing data from the 1000 Genomes Project, teaching it CNV-calling capabilities similar to conventional methods. Next, the model is fine-tuned using high-confidence CNV calls derived from only a few existing high-coverage aDNA samples. During this stage, the model adapts to making CNV calls based on the downsampled read depth signals of the same aDNA samples. LYCEUM achieves accurate detection of CNVs even in typically low-coverage ancient genomes. We also observe that the segmental deletion calls made by LYCEUM show correlation with the demographic history of the samples and exhibit patterns of negative selection inline with natural selection. LYCEUM is available at https://github.com/ciceklab/LYCEUM.

## 1 Introduction

The analysis of ancient DNA (aDNA) offers valuable insights into species evolution, human migration, and historical population dynamics. With advances in next-generation sequencing technologies, aDNA studies have evolved to reconstruct entire genomes, trace disease spread, and explore human ancestry, thus providing a more comprehensive understanding of genetic variation over time [1]. Despite these advancements, much of the genetic variation analysis in aDNA samples has predominantly focused on single nucleotide polymorphisms (SNPs), while the analysis of Copy Number Variations (CNVs) remains equally critical for understanding phenotypic diversity.

Gene duplication and deletion variants often exert a more substantial impact on phenotypes compared to point mutations and are key drivers of phenotypic variation that allow species to adapt to their environments [2–6]. For example, in human evolution, it has been shown that the expansion of the human-specific copy number of TCAF genes occurred approximately 1.7 million years ago, likely facilitating adaptation to cold climates or dietary changes [7]. Furthermore, the TBC1D3 gene family, which plays a role in modulating epidermal growth factor receptor (EGFR) signaling, underwent significant expansion after the divergence of humans from chimpanzees [8]. This expansion is associated with the development of the human prefrontal cortex, which highlights the potential contribution of CNVs to the evolution of the human brain. Together, these findings underscore the pivotal role of CNVs in human evolutionary processes. Furthermore, analysis of CNV in aDNA samples offers the potential to illuminate the historical prevalence of risk factors for complex diseases associated with CNV. Recent research has highlighted the contribution of CNVs to susceptibility to complex traits and conditions, such as autism spectrum disorder [9, 10], schizophrenia [11], and bipolar disorder [12]. A recent study of genome-wide association of CNV (GWAS) identified 73 signals associated with 40 different diseases [13], all of which increase the risk of their respective conditions. Despite the well-documented role of CNVs in disease susceptibility, limited research has focused on the origins of these variants. Thus, examining CNVs between diverse ancestral groups may offer insight into the origins of common disease susceptibilities in modern populations.

However, detecting structural variants (SVs), including CNVs, in aDNA samples poses considerable challenges. aDNA typically exhibits extensive postmortem damage, including deamination and fragmentation [14]. In addition, pervasive microbial contamination often results in low endogenous DNA content, further complicated by contamination by closely related species, which makes it difficult to distinguish endogenous DNA from contaminant sequences [15, 16]. As a result, endogenous DNA in aDNA samples often has low sequencing coverage, which complicates the discovery of SVs. State-of-the-art whole genome sequencing (WGS) and whole exome sequencing (WES)-based CNV callers rely on higher coverage [17–21, **?**,**?**,**?**] and long-read sequencing data [22, 23], making these approaches suboptimal for low-coverage aDNA samples. CONGA is the only method specifically designed for the detection of CNVs in aDNA [24]. It uses GC-content normalization to account for sequencing biases and employs a likelihood-based scoring system, termed the C-score, to assess the probability of CNV presence. However, the performance is limited especially in low coverages due to the high noise levels and complexity of the problem.

To address these challenges, we introduce LYCEUM, the first deep learning-based approach to detect CNVs in low-coverage and contaminated aDNA samples. LYCEUM processes the read depth signal of each exon using a cascade of convolution and transformer encoder blocks which also incorporates a trainable classification token. This token is chromosome specific and is used to determine if a CNV event has occurred within the exon. The model captures the chromosome-specific context using this token. However, the scarcity of high-quality aDNA samples bar training such a deep and complex model, which is why deep learning has not been employed in this domain yet. To overcome this, we first pre-train the model using CNV calls obtained for hundreds of WGS samples from the 1000 Genomes project [25], using conventional CNV callers (e.g., DRAGEN [26] or CNVnator [18]). This way, the model learns the relationship between the high-quality read-depth signal and CNV events on the WGS data. To adapt the model using aDNA samples, we fine-tune it using a few available high-coverage aDNA samples and some simulated aDNA samples. That is, we fine-tune the model with (i) the reliable CNV calls obtained from high-coverage aDNA read depth signal using again the conventional callers as labels, and (ii) the down-sampled versions of the corresponding read depth signal as features, to mimic typically very-low-coverage aDNA samples. This novel approach teaches the pre-trained model to relate the low-coverage read depth signal with the CNV events. Extensive benchmarks show that LYCEUM consistently achieves the state-of-the-art CNV calling performance for a wide range of coverage levels and works well with even very low coverages (e.g., 0.05*x*). We also observe that the segmental deletion calls identified by LYCEUM are closely correlated with the demographic history of the samples and display patterns of negative selection due to natural selection. Both confirm the reliability of the calls made by LYCEUM. Given its robustness in CNV calling and its adaptability for aDNA samples through fine-tuning, we think LYCEUM will have a big impact on CNV detection and genomic analyses of low-coverage ancient datasets in the fields of archaeogenetics and paleogenomics. LYCEUM is available at https://github.com/ciceklab/LYCEUM.

## 2 Methods

### 2.1 Problem Formulation

LYCEUM detects CNVs in the coding regions. Let *X* be the set of all exons in the genome and let *X*^*i*^ indicate the *i*^*th*^ exon where *i* ∈ *{*1, 2, …., *N*} and *N* = |*X*|. Each *X*^*i*^ is associated with (i) a chromosome 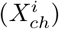 where *ch* ∈ *{*1, 2, 3, …, 24*}*. 23 and 24 represent chromosomes X and Y, respectively; (ii) coordinates within that chromosome: 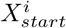 and 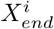 and finally, (iii) a normalized read depth signal vector: 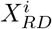 which can be at most 1 kbp-long and padded from right, if shorter. Longer exons are discarded, which is relatively rare. *Y* ^*i*^ is the semi-ground truth label for *X*^*i*^, obtained using the conventional CNV caller tools such as CNVnator or DRAGEN on the high-coverage read-depth signals of all aDNA samples. Let *Ŷ*^*i*^ = *f* (*X*^*i*^, *θ*) be the CNV prediction (deletion, duplication, or no-call) using the model *f* (a multi-class classifier) which is parameterized by *θ*. The goal is to find the model parameters *θ* that minimize the difference between the predicted exon-level CNV labels and their ground truth labels.

### 2.2 Model Description

We show the architecture of LYCEUM in Figure 1. The model first inputs the read depth signal 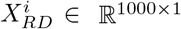 and the signal is embedded by two sequential 1D convolutional layers. The first layer employs 32 filters with an input channel size of 1 and an output channel size of 32, and the second layer employs 64 filters with an input channel size of 32 and an output channel size of 64, both using a kernel of size 3 and stride 1. Each convolution layer is followed by the Batch Normalization (BN) and the ReLU activation function. It results in an embedded vector 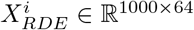. A trainable classification token *c*^*t*^ ∈ ℝ^64^, which is specific to each chromosome *t*, is appended to 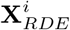 to obtain the final input tensor 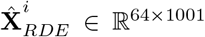 which will be input into the cascade of transformer encoder blocks. This classification token contains the learnable parameters of the model and is used to classify the label for this exon in the context of this chromosome later in the model and also informs the model about the context of other exons in this chromosome. Finally, 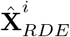 is summed with a same-sized positional encoding tensor to inform the model about the exact position of each read depth value embedding within this exon. This tensor is passed through a cascade of three transformer encoding blocks to obtain the transformed output of the same size 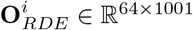. The last column of the tensor that contains the chromosome-specific token is sliced 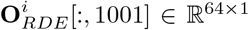 and passed through a fully connected neural network followed by softmax to predict the probabilities of DEL, DUP and NO-CALL for exon *X*^*i*^. *Ŷ*^*i*^ is assigned as the class that has the highest probability among these three options.

**Fig. 1:**
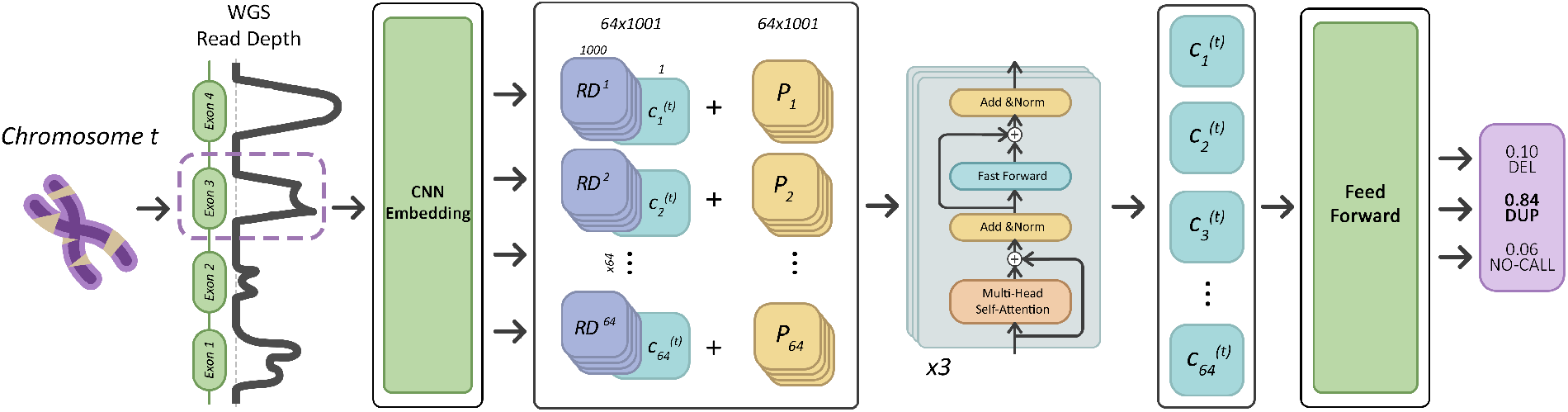
The system model of LYCEUM. The read depth (RD) for each exon (1kbp) is embedded using 2 sequential convolution layers with 32 and 64 kernels, respectively. Batch normalization (BN) and the rectified linear unit (ReLU) activation function are applied after each convolutional operation. The resulting vector (64×1000) concatenated with a chromosome-specific classification token (c) (64×1) and summed up with positional encoding (P, 64×1001). This tensor is passed through 3 transformer encoder blocks and the sliced classification token is input to a fully connected neural network which outputs the call probabilities.

**Fig. 2:**
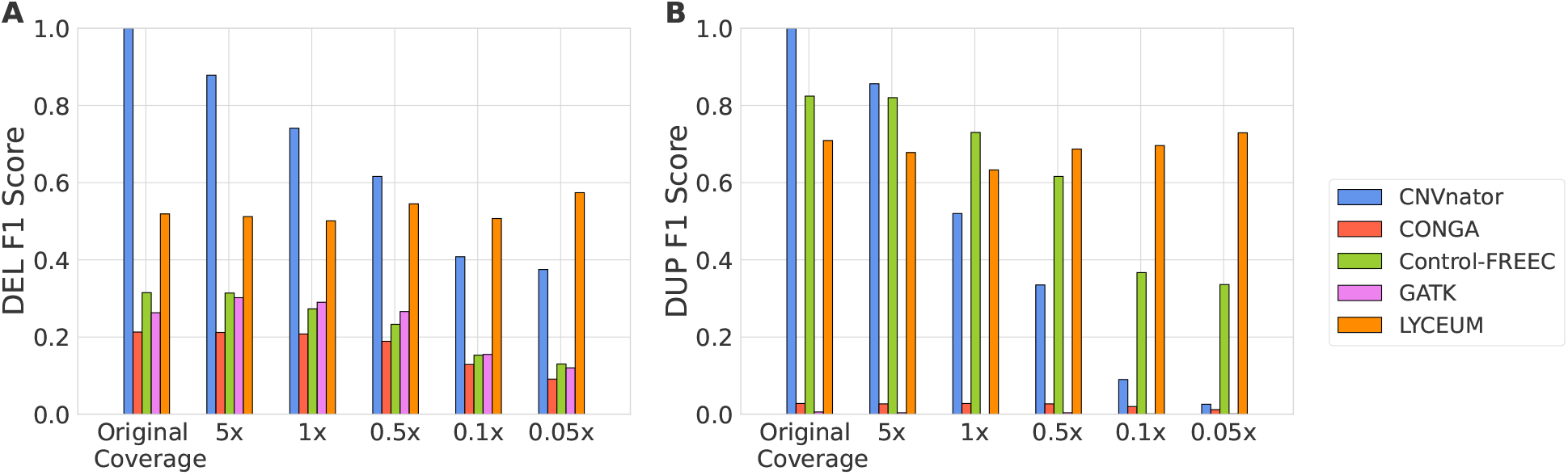
A comparative analysis of F1 scores for CNV caller algorithms applied to seven ancient genome samples with varying coverage levels across gene regions. CNVnator calls on the original coverage to serve as a semi-ground truth reference. Each sample is down-sampled to five different coverage levels, and each algorithm is subsequently tested on these samples. **(A)** F1 score results for deletion detection. **(B)** F1 score results for duplication detection.

Let *LYCEUM* (*X*^*i*^, *θ*) be the LYCEUM model which is the multi-class classifier *f* that performs exon-wise CNV prediction (ie DEL, DUP, NO-CALL) using the WGS-based read-depth signal, parameterized by *θ* that is pre-trained using the high coverage WGS samples and DRAGEN calls obtained from the samples from the 1000 Genomes dataset. This model is fine-tuned using a few available aDNA WGS samples, which have high coverage, to obtain *θ*^*′*^. The features used are various down-sampled versions of the read depth signal for aDNA samples. The parameters of the model *θ*^*′*^ are optimized to minimize the difference between *Ŷ*^*i*^ and the semi-ground truth labels *Y* ^*i*^ obtained using CNVnator on the original high coverage version of the read depth signal. *Ŷ*^*i*^ = *LY CEUM* (*X*^*i*^, *θ*^*′*^) is the prediction of the LYCEUM model for the exon *X*^*i*^.

Approximately 5% of ancient exon targets lack the read depth information which prohibits LYCEUM to make any calls for those exons. We address this issue by applying majority voting for that exon based on *Ŷ*^*I*^ of the three nearest-neighbor exons. To obtain gene-level calls, LYCEUM performs majority voting across all exons of that gene.

### 2.3 Data Sets

#### Pretraining Dataset

We use 550 randomly selected WGS samples from the 1000 Genomes Project to pretrain the model. The IDs for these samples are provided in the Supplementary Table 28. We have DRAGEN-based CNV calls available for these samples through *1000 Genomes Phase 3 Reanalysis with DRAGEN 3*.*5, 3*.*7, 4*.*0, and 4*.*2* which was accessed on Jan 2025 from https://registry.opendata.aws/ilmndragen-1kgphttps://registry.opendata.aws/ilmn-dragen-1kgp.

#### Ancient Genome Dataset

We use 63 WGS samples aDNA from west and east Eurasia, as well as North America [27–45]. We present the metadata for these datasets in Supplementary Tables 25-27. The levels of coverage of the samples vary widely, ranging from 0.04 to 26.3 (median = 4.2). To obtain reliable ground truth, we separate 20 moderate coverage samples (>9x) and run CNVnator on these samples. We obtain 31, 946 DUP and 55, 337 DEL calls in total at the exon level and 3, 324 DUP and 6, 823 DEL calls at the gene level. True negative samples are the remaining exons or genes for which CNVnator has not made any calls for exon-specific and gene-specific settings, respectively. These calls collectively form the semi-ground truth CNV calls. Then, we down-sample the read depth of each sample into 5x, 1x, 0.5x, 0.1x and 0.05x coverage versions. Among the 20 samples, we randomly choose 7 samples (together with the corresponding downsampled versions) in the data set and assign them to the test set. We use the samples with IDSs “Ust", “VLASA7", “Bichon", “VLASA32", “AKT16", “Nea3", and “STAR1” as test samples, and the rest of the dataset (13 samples) as training samples. The test samples are used to evaluate all models (see Supplementary Tables 25-26 for details). Specifically, the training samples in the dataset are labeled with 18, 967 DUP and 47, 795 DEL calls in total for the exon-specific setting and, with 2, 019 DUP and 5, 730 DEL calls for the gene-specific setting. On the other hand, test samples in the dataset are labeled with 12, 979 DUP and 7, 542 DEL calls for the exon-specific setting and, with 1, 305 DUP and 1, 093 DEL calls for the gene-specific setting.

For the analysis of segmental deletions, which is discussed in Sections 3.5 and 3.6, we use the remaining 43 samples combined with the 7 test samples mentioned above (See Supplementary 27 for details). The coverage values of these 50 samples are between 0.04 and 26.3 (median = 3.5). Note that we do not include down-sampled versions of these 7 samples in these analyses.

#### Simulated Ancient Genome Dataset

We use the VarSim [46] tool to insert DEL and DUP variations into the reference genome hg38. This method is a variant simulation framework used for both germline and somatic mutations. We randomly select 15 WGS samples from the 1000 Genomes Project. We then input these 15 labels to the VarSim tool with the following parameters: sv_num_del = 2000, sv_num_dup = 2000, sv_percent_novel = 0.01, sv_min_length_lim = 500, sv_max_length_lim = 100000 and total_coverage = 10. These parameters allow us to simulate novel DEL and DUP variations while preserving a genome’s variational distribution. The simulated 15 genomes contain DEL and DUP insertions in the size range of 500 bps and 100 kbps. Next, we use the Gargammel [14] tool to simulate ancient genomes. This tool was specifically developed to generate artificial ancient genome fragments. We input the 15 simulated genomes with DEL and DUP insertions into Gargammel with the following parameters: nick frequency = 0.024, length of overhanging ends = 0.36, probability of deamination of Cs in double-stranded parts = 0.0097, and probability of deamination of Cs in single-stranded parts = 0.55. These parameters allow us to simulate postmortem DNA damage, including fragmenting DNA into shorter sequences and adding degradation patterns typical of aDNA samples. We run Gargammel with coverages of 1x, 0.5x, 0.1x, and 0.05x for each Varsim output, in order to further mimic low-coverage aDNA. We randomly pick 5 Varsim generated genomes and use their corresponding 20 Gargammel-processed versions for training, and we use the remaining 40 samples as test samples. The samples in the test set do not overlap with the preraining dataset. All of the 60 samples in the dataset are labeled with 43,335 DUP and 21,469 DEL calls for the exon-specific setting, and with 7,313 DUP and 4,459 DEL calls for the gene-specific setting. True negative samples are the remaining exons or genes for which CNVnator has not made any calls for exon-specific and gene-specific settings, respectively.

## 3 Results

### 3.1 Experimental Setup

#### Compared Methods

We compare LYCEUM with the following WGS-based CNV callers: CNVnator[18], CONGA[19], Control-FREEC[24], and GATK[47]. We use CNVnator’s CNV calls on full-coverage data as semi-ground-truth labels and test CNVnator and others on down-sampled real aDNA data and the simulated aDNA data. LYCEUM, CNVnator, and CONGA report categorical CNV predictions. Control-FREEC reports integer CNVs. To include this method in our comparison, we discretize its predictions. We use DEL if the tool’s output is less than 2, DUP if it is larger than 2 and NO-CALL if it is 2. GATK results are presented only for real aDNA samples. It consistently fails to process simulated aDNA samples.

#### Parameter Settings

For CNVnator, we use the recommended bin size parameter based on coverage in their paper. Specifically, we use bin size 1,000 for full coverage, 3,000 for 5x, 15000 for 1x, 30,000 for 0.5x, 150,000 for 0.1x and 300,000 for 0.05x data. For Control-FREEC, all parameters are set to their default values. For CONGA, we evaluated various configurations with minimum c-values [0.3, 0.5] and min-sv-size values [100, 1000]; yet, the default parameters were ultimately selected as they provided better performance.

#### Training LYCEUM

We pre-trained our model using 550 randomly selected WGS samples from the 1000 Genomes Project for one epoch using the DRAGEN-based CNV event calls. Then, we fine-tuned it using (i) WGS-based 13 aDNA training samples; (ii) Five down-sampled versions of each of the samples in (i) (5x, 1x, 0.5x, 0.1x, 0.05x), and 20 simulated aDNA samples at 4 coverage levels (1x, 0.5x, 0.1x, 0.05x). The read depth signals overlapping with exon regions are used as input, with CNV calls from the original full-coverage aDNA data generated by CNVnator serving as labels for real samples, while the output VCF file from VarSim provides the ground truth for simulated samples. We name this model LYCEUM. During fine-tuning, we use Adam optimizer [48] and cosine annealing learning rate scheduler with an initial learning rate of 5*x*10^−5^. The model converged in 3 epochs.

We train all models on a SuperMicro SuperServer 4029GP-TRT with 2 Intel Xeon Gold 6140 Processors (2.3GHz, 24.75M cache) and 256GB RAM. We used a single NVIDIA GeForce RTX 2080 Ti GPU (24GB, 384Bit). Fine-tuning took approximately 72 hours. The average time to call all CNVs per ancient genome is ~ 5 mins.

### 3.2 Evaluation criteria

LYCEUM classifies each exon as deletion, duplication, or no-call. So we report exon-level performance (precision, recall, and F1 score). For comparison, we map CNV calls from the compared methods to the target exon regions and label each corresponding exon accordingly. In addition to this, we also report gene-level performance. To determine the overall CNV label for a gene, we aggregate the predicted CNV labels of all the corresponding exons and assign a final gene-level CNV label using majority voting. This means that the most frequent CNV label (deletion or duplication) among exons is selected as the final gene-level CNV label. The predicted CNV labels for exons of a gene are mostly consistent and the cases where both deletion and duplication labels are called for exons within the same gene are rare (0.2%). To ensure a fair comparison of performance across different models, we standardize this process by applying the same majority voting methodology to all models being compared. First, the predicted CNV calls from each model were mapped to the target exon regions. After aligning the exon-level predictions, we determine the gene-level CNV label for each model using the same majority voting strategy.

### 3.3 The performance of LYCEUM on low-coverage aDNA samples

To evaluate the performance of LYCEUM in low-coverage aDNA samples, we use 20 aDNA samples for which we have moderate coverage (>9x) obtained from eight published studies [27–34]. Since these samples have relatively higher coverage compared to a typical aDNA which has <1x coverage, we treat them as regular WGS samples and use the WGS CNV caller CNVnator [18] to obtain semi-ground truth CNV calls. To assess the robustness of the models at varying depths of sequencing coverage, we downsample samples to specific coverage levels, 5x, 1x, 0.5x, 0.1x, and 0.05x, using samtools-view [49]. This step allows us to simulate the challenges posed by low-coverage ancient genomes, and observe how well the models maintain performance under such conditions. We use 13 samples in the training set and test the model on the remaining 7 samples. For benchmarking, we compare LYCEUM against the only available aDNA CNV caller CONGA and the state-of-the-art WGS CNV callers CONTROL-FREEC, GATK and CNVnator. Note that there is no groundtruth CNV call set for aDNA samples. Our goal with LYCEUM is to be able to call CNVs on low-coverage aDNA data with performance similar to that of CNV calling on moderate-to-high-coverage WGS data.

Figure 2 shows the comparison of the F1 scores of the methods to detect deletions and duplications in genes. We also present the exon-level and gene-level precision and recall results in Supplementary Figures 1 and 2, respectively, along with the corresponding confusion matrices in Supplementary Tables 3-12. Note that CNVnator’s F1 score on the original coverage is 1.0 since its predictions are used as the semi-ground truth calls to measure performance. Yet, as the coverage goes down, its performance consistently deteriorates for CNVnator. CONGA achieves very high precision but very low recall using the suggested hyperparameters (see Supplementary Tables 3-12) and its performance also deteriorates with lower coverage. Note that these are the hyperparameters that performed the best in our hyperparameter optimization. On the other hand, LYCEUM demonstrates consistent performance across all coverage levels for deletions and duplications, maintaining its performance even at very low coverages (0.1x and 0.05x), where it outperforms other tools. We also observe that it maintains higher recall as opposed to CONGA, close to the level of a regular WGS-based CNV caller. It achieves a 20% and 39% F1 score improvement over the next-best method at the lowest coverage (0.05x) for deletions and duplications, respectively. We also measure the contamination rates of test samples using ContaminationX [50] and observe that high contamination rate does not affect the performance of LYCEUM (See Supplementary Table 23). This highlights LYCEUM’s robustness in low-coverage and highly contaminated conditions, a critical advantage in ancient genome analysis, where such scenarios are common.

### 3.4 Comparative performance analysis on simulated ancient genomic data

Figure 3 compares the results of the methods on simulated aDNA data as explained in Section 2.3. Detailed exon-level and gene-level performance scores are provided in Supplementary Tables 13, and 14, along with corresponding confusion matrices in Supplementary Tables 15-22. We observe that LYCEUM exhibits superior performance compared to other methods at the lowest coverage value (0.05x), achieving F1 score improvements of 60% and 68% over the second-best method for deletions and duplications, respectively.

**Fig. 3:**
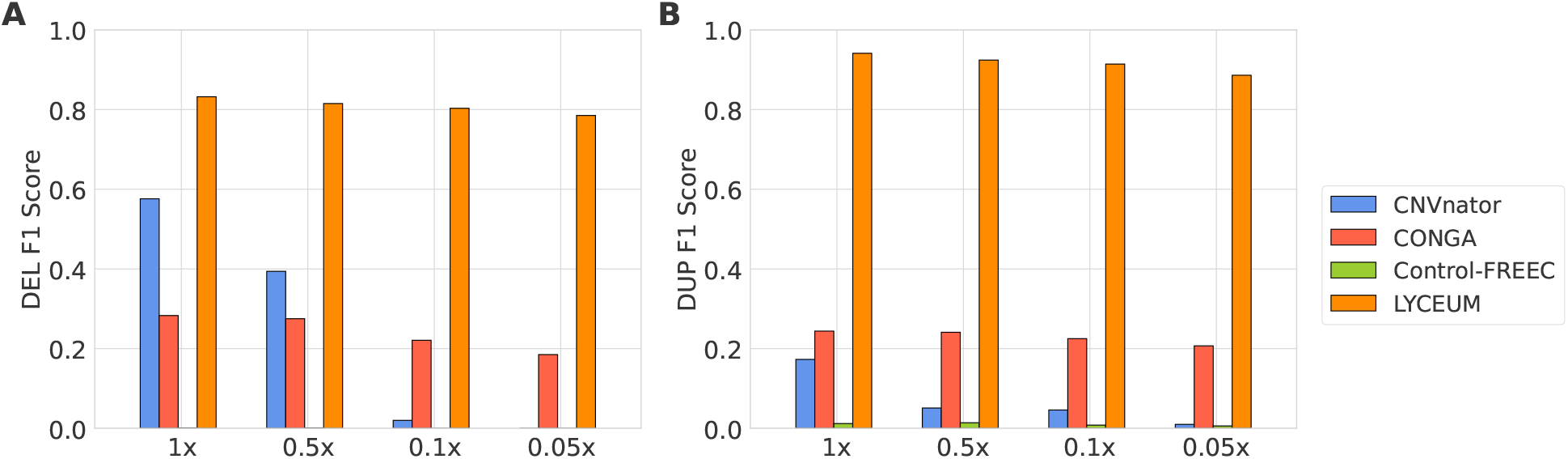
Comparison of F1 scores among CNV caller algorithms evaluated on simulated ancient genomes with varying coverage levels in gene regions. **(A)** F1 score performance for detecting deletions. **(B)** F1 score performance for detecting duplications. GATK was excluded from the simulated ancient genome analysis because it consistently failed to process the simulated samples due to recurring exceptions.

Furthermore, LYCEUM’s performance remains highly consistent with decreasing coverage, highlighting its robustness. This suggests that the LYCEUM’s architecture is particularly effective for scenarios involving highly degraded and incomplete datasets, where conventional tools often struggle.

### 3.5 Segmental deletions detected by LYCEUM recover the demographic history

To assess the reliability of LYCEUM, we analyze the distribution of high-confidence (probability > 0.9) and variable (variance > 0.25) exon-level deletion calls across 50 aDNA samples. The geographic locations of the findings are shown in Figure 4A where darker blue indicates western Europe and darker red indicates eastern Asia. We apply Principal Component Analysis (PCA) to visualize the genetic background of these samples in a 2D space, shown in Figure 4B. In particular, the PCA plot reveals two primary clusters, corresponding to samples from Europe and Asia, that align with the geographic distribution. We observe that samples mostly cluster among west and east origins and the samples from central Asia are distributed across two clusters. Furthermore, we compare the clustering performance of LYCEUM with CONGA. Figure 4C presents the PCA plot of the deletion events detected by CONGA with variance > 0.25, across the same 50 aDNA samples. The subsequent calculation of the silhouette index reveals that LYCEUM achieves a silhouette index of 0.12, while CONGA has a slightly lower value of 0.09. Additionally, hierarchical clustering analysis, as shown in Figure 5, indicates a clustering accuracy of 70% for LYCEUM and 66% for CONGA. These results indicate that both LYCEUM and CONGA detect CNVs that are concordant with demographics, while LYCEUM is performing slightly better.

**Fig. 4:**
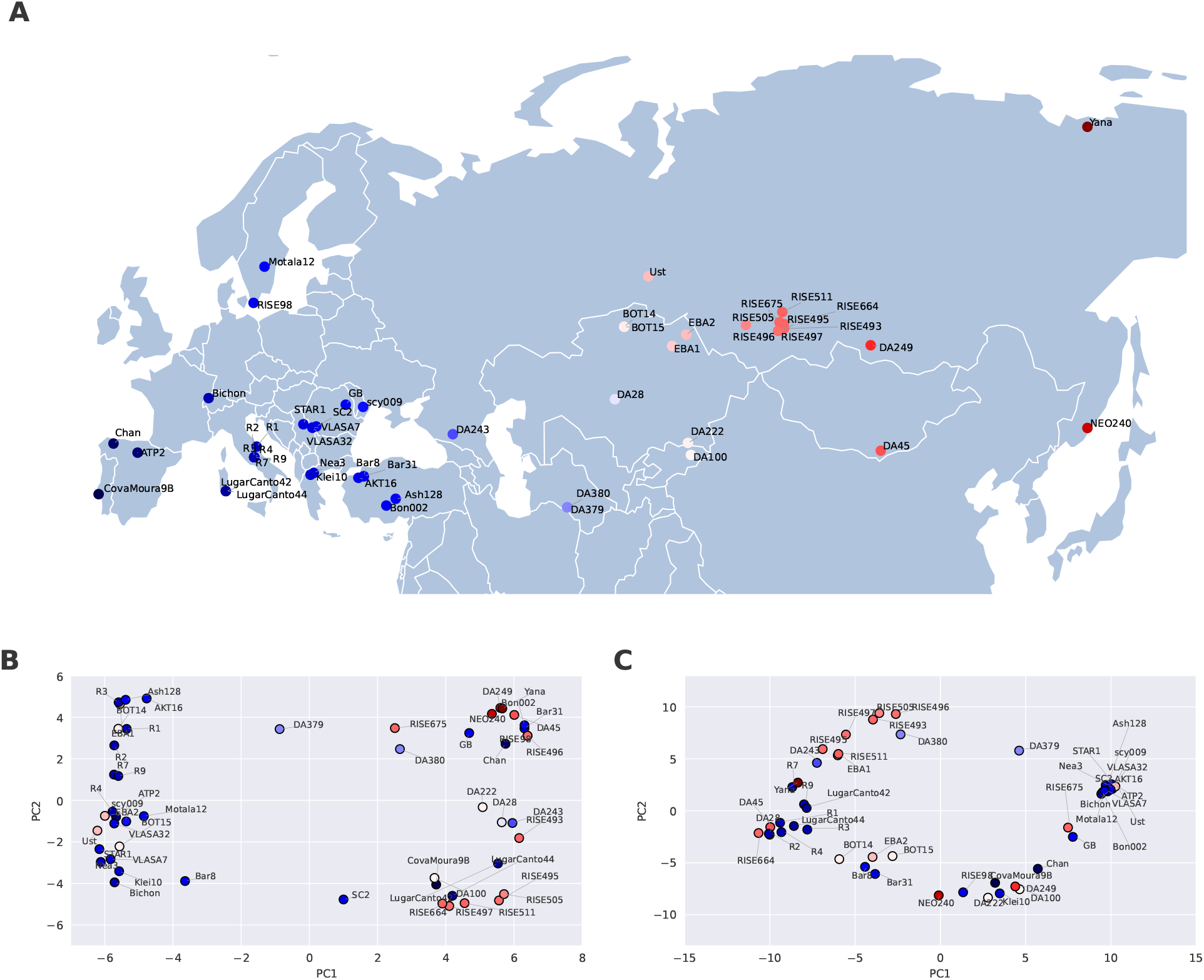
Distribution of segmental deletions (**A**) Geographic locations of 50 aDNA samples. The color spectrum represents the relative positions of samples from east (red) to west (blue). (**B**) and (**C**) show the visualization of the first two principal components representing genomic distances based on segmental deletion events identified by LYCEUM and CONGA, respectively. Results highly correlate with demographic structure

**Fig. 5:**
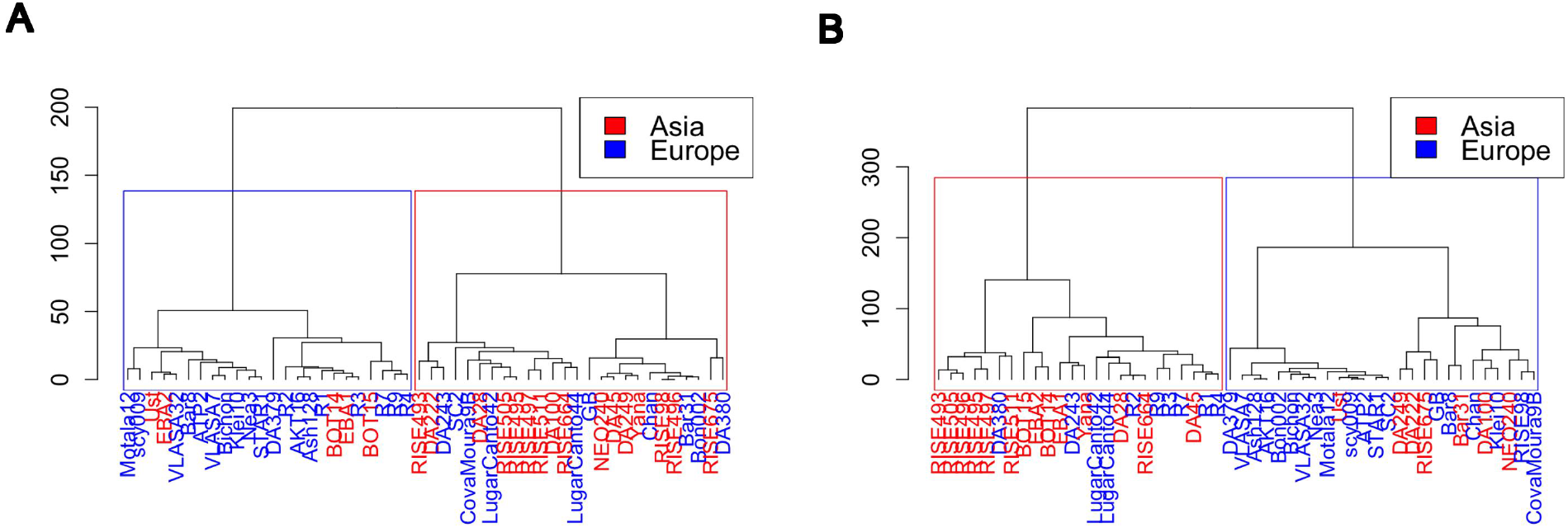
Hierarchical clustering dendrograms of LYCEUM (**A**) and CONGA (**B**) on 50 aDNA samples. Both dendrograms demonstrate a similar clustering trend according to geographic origin, highlighting genetic distinctions between ancient European and Asian populations.

### 3.6 Segmental deletions are negatively selected

Deletion events often negatively impact fitness. These deletion variants are subject to negative selection and are gradually eliminated from the gene pool over time [51, 52]. Moreover, longer deletions are observed less frequently due to stronger negative selection, likely because they have more significant functional consequences [53]. In our analysis shown in Figure 6 we use 89,964 autosomal deletion events detected by LYCEUM on 50 aDNA samples to prove that the calls for LYCEUM are biologically consistent and align with known evolutionary dynamics.

**Fig. 6:**
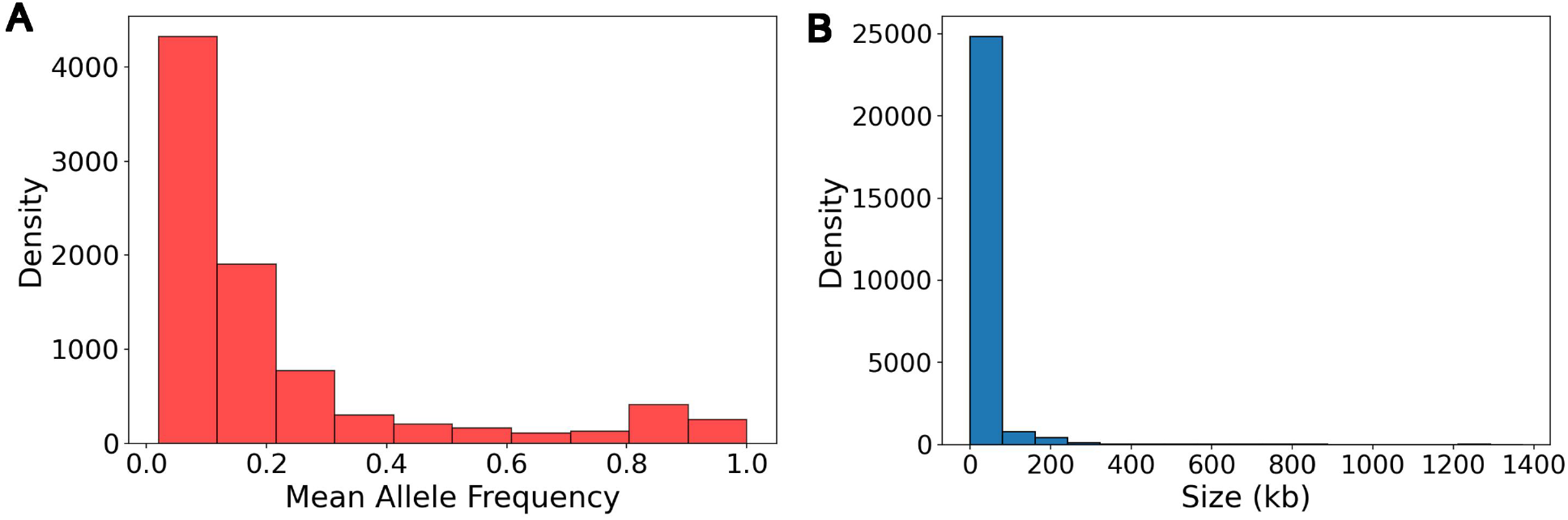
Impact of negative selection autosomal deletion variants. **(A)** Analysis of the relationship between the mean allele frequency of these deletions and their density, illustrating how allele frequency influences the prevalence of deletion variants across the genome. **(B)** Examination of the relationship between deletion size and the density of these deletions, providing insights into how the size of deletion variants correlates with their distribution frequency within autosomal regions.

The Site Frequency Spectrum (SFS) in Figure 6A illustrates the relationship between the mean allele frequency and the density of deletions. The figure shows that deletions with lower allele frequencies have higher densities, indicating that less frequent deletions are much more dense in the autosomal deletion call set. This pattern is consistent with the expected influence of negative selection, where rare deletions are maintained at low frequencies due to selective pressure against them [51]. Moreover, Figure 6A shows that 74% of the detected positions have allele frequencies below 20%, demonstrating that the model is not memorizing high-frequency events but is effectively identifying rare CNVs rather than overfitting to commonly observed variants.

The results in Figure 6B indicate that longer deletions are substantially less frequent. This observation is consistent with the expectation that longer deletions face stronger negative selection pressures [53]. Specifically, we found a strong negative correlation (Spearman correlation = −0.88) between deletion size and density. It shows that natural selection acts more intensively against larger deletions, leading to their lower representation in the population and our predictions are inline with this fact.

## 4 Discussion

The results of this study reveal the efficacy of LYCEUM, our deep learning-based CNV caller, in addressing the unique challenges associated with aDNA analysis, particularly in detecting CNVs from low-coverage and noisy genome data. As demonstrated, LYCEUM maintains high performance in varying depths of sequencing coverage, substantially outperforming other CNV callers such as CONTROL-FREEC, GATK, CNVnator, and also the state-of-the-art ancient genome CNV caller CONGA, especially in very low coverage (0.05x). The method’s performance is consistent across coverage levels. We tested the limits of this approach and investigated the coverage levels where the method is no longer effective. We found that only at ultra low coverage levels such as ≤ 0.01x, the method’s performance declines where the read depth signal is no longer informative (See Supplementary Figures 1-2). We also show that the detected CNV composition can accurately recover the geographical origins of the samples and reconfirm that deletion events are under negative selection.

While our test and training (fine-tuning) datasets are distinct, one possible concern is that the fixated CNVs in the population might overlap between the two sets. This might leaks information. Our analysis reveals that 63% of duplications and only 18% of deletions are shared between the training and test sets. This indicates that duplications show higher overlap, and deletions remain largely distinct. While the DUP calling performance is slightly higher on average which might be due to the higher overlap, DEL calling performance is also very high and substantially outperforms the state-of-the-art. This shows that LYCEUM’s performance improvement does not stem from information leak but because of learning the relation between the read depth signal and the CNV events better. We also ensure that the 1000 Genomes samples used for simulating aDNA samples and testing the performance do not overlap with the 1000 Genomes samples used for pretraining to prevent information leakage.

Despite these promising results, some limitations remain. The reliance on CNVnator or DRAGEN calls as semi-ground truth for real aDNA samples, while necessary due to the lack of true ground truth, might be introducing potential biases into performance evaluation. Future work will aim to validate LYCEUM’s predictions using independently verified CNV datasets from ancient genomes as they become available.

Using DRAGEN calls as labels for pretraining, and CNVnator calls as labels for fine-tuning potentially limits the performance of the algorithm as these tools might disagree on the labels which would make the training of LYCEUM more challenging. We made this choice because DRAGEN calls were freely available for a large sample set. Although we present results in this more challenging setting, we obtain state-of-the-art results which show the versatility of the method but also suggest room for improvement for LYCEUM.

Currently, the method works on coding regions because these represent the primary focus to understand the history of genetic diseases and we need to limit the scope of the input for the model training to scale. Future work will first focus on incorporating important regulatory non-coding regions, such as evolutionarily conserved regions, into the training process and then to scale to the full genome. Integrating these regions may broaden the scope of detected CNVs and provide deeper insights into their functional implications within ancient genomes. We also note that the model can easily be extended by the users to incorporate regions-of-interest by defining those region(s) as mock exon(s) to be input to the model.

In conclusion, LYCEUM represents a significant advance in the field of aDNA analysis, offering a powerful tool for CNV detection in low-coverage and highly degraded datasets. Its ability to maintain accuracy across different coverage levels, combined with its scalability through a fine-tuning strategy, makes LYCEUM a valuable resource for researchers studying evolutionary processes, human migration, and the genetic underpinnings of disease susceptibility in ancient populations.

## Data Availability

LYCEUM is implemented and released at https://github.com/ciceklab/LYCEUM under CC-BY-NC-ND 4.0 International license. Samples, semi-ground-truth-labels, and CNV calls made by LYCEUM can be accessed through https://zenodo.org/records/13997011 to reproduce these analyses.

## Acknowledgments

We would like thank Alperen Utku Yalçın, Erfan Farhang Kia, Dilek Koptekin, Can Alkan and Mehmet Somel for their valuable contributions to this paper.

