## Supplementary Material for "LYCEUM: Learning to call copy number variants on low coverage ancient genomes"

### Supplementary Material
### for
### LYCEUM: Learning to call copy number variants on low coverage ancient genomes

#### 1 Supplementary Figures

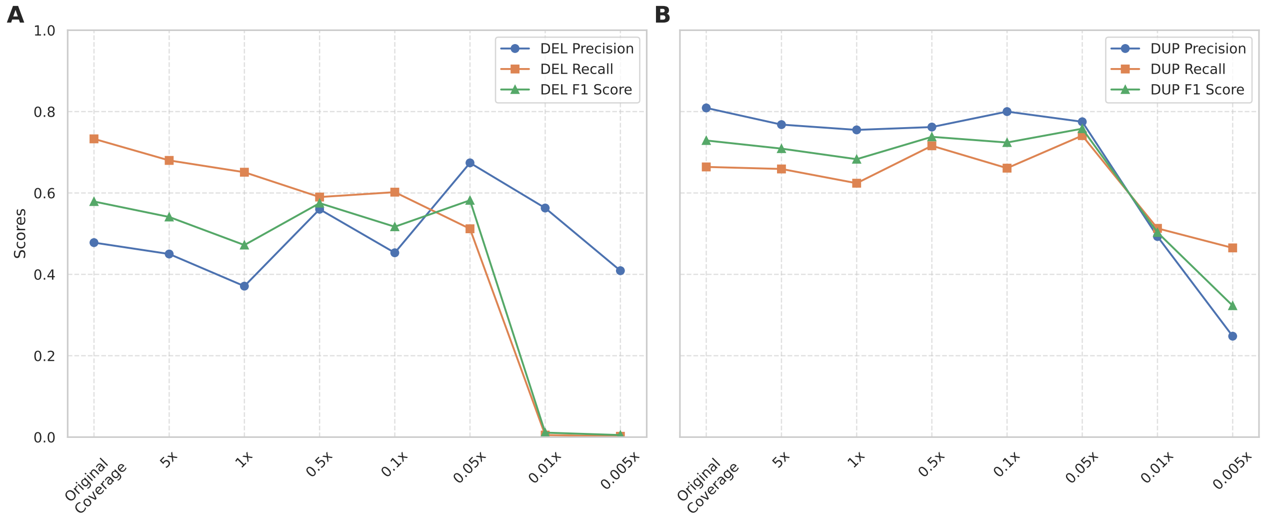


**Supplementary Figure 1.** Comparison of LYCEUM's performance in detecting deletion (A) and duplication (B) events in test samples across varying coverage levels, including low and ultra-low coverages in exon regions.

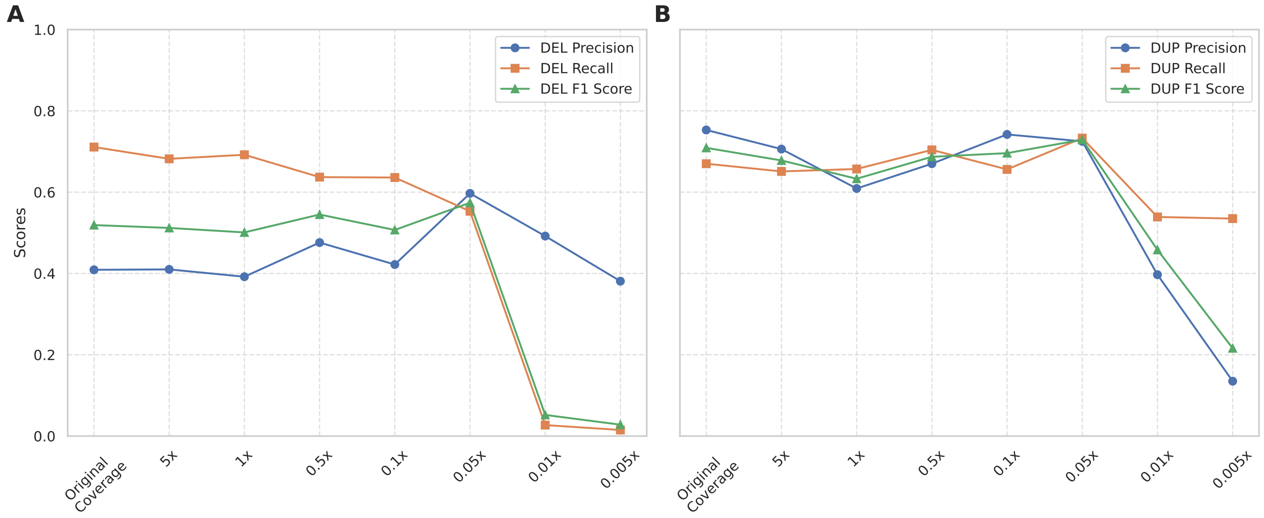


**Supplementary Figure 2.** Comparison of LYCEUM's performance in detecting deletion (A) and duplication (B) events in test samples across varying coverage levels, including low and ultra-low coverages in gene regions.

#### 2 Supplementary Tables

**Supplementary Table 1.** The performance comparison of the CNV callers on the down-sampled ancient samples (test set) in exon regions. CNVnator calls on original coverage samples are used as the ground truth.

| Coverage | Tools | DEL Precision | DEL Recall | DEL F1 Score | DUP Precision | DUP Recall | DUP F1 Score | Overall Precision | Overall Recall | Overall F1 Score |
|---|---|---|---|---|---|---|---|---|---|---|
| Original Coverage | CNVnator | 1.0 | 1.0 | 1.0 | 1.0 | 1.0 | 1.0 | 1.0 | 1.0 | 1.0 |
| | CONGA | 0.838 | 0.093 | 0.167 | 0.528 | 0.009 | 0.017 | 0.683 | 0.051 | 0.092 |
| | Control-FREEC | 0.176 | 0.826 | 0.291 | 0.861 | 0.814 | 0.837 | 0.5185 | 0.82 | 0.564 |
| | GATK | 0.133 | 0.89 | 0.231 | 0.233 | 0.013 | 0.025 | 0.183 | 0.4515 | 0.128 |
| | LYCEUM | 0.478 | 0.733 | 0.579 | 0.809 | 0.664 | 0.729 | 0.644 | 0.699 | 0.654 |
| 5x | CNVnator | 0.928 | 0.913 | 0.92 | 0.914 | 0.826 | 0.868 | 0.921 | 0.87 | 0.894 |
| | CONGA | 0.835 | 0.094 | 0.169 | 0.533 | 0.008 | 0.016 | 0.684 | 0.051 | 0.0925 |
| | Control-FREEC | 0.176 | 0.829 | 0.29 | 0.889 | 0.804 | 0.845 | 0.5325 | 0.8165 | 0.5675 |
| | GATK | 0.15 | 0.588 | 0.239 | 0.091 | 0.005 | 0.009 | 0.1205 | 0.2965 | 0.124 |
| | LYCEUM | 0.45 | 0.68 | 0.541 | 0.768 | 0.659 | 0.709 | 0.609 | 0.67 | 0.625 |
| 1x | CNVnator | 0.843 | 0.779 | 0.81 | 0.879 | 0.464 | 0.607 | 0.861 | 0.622 | 0.709 |
| | CONGA | 0.769 | 0.097 | 0.173 | 0.504 | 0.009 | 0.017 | 0.6365 | 0.053 | 0.095 |
| | Control-FREEC | 0.163 | 0.765 | 0.268 | 0.845 | 0.694 | 0.762 | 0.504 | 0.7295 | 0.515 |
| | GATK | 0.145 | 0.554 | 0.23 | 0.009 | 0.002 | 0.004 | 0.077 | 0.278 | 0.117 |
| | LYCEUM | 0.371 | 0.651 | 0.472 | 0.755 | 0.624 | 0.683 | 0.563 | 0.638 | 0.578 |
| 0.5x | CNVnator | 0.785 | 0.671 | 0.723 | 0.76 | 0.252 | 0.378 | 0.773 | 0.462 | 0.551 |
| | CONGA | 0.639 | 0.097 | 0.168 | 0.479 | 0.008 | 0.016 | 0.559 | 0.0525 | 0.092 |
| | Control-FREEC | 0.143 | 0.655 | 0.235 | 0.828 | 0.593 | 0.691 | 0.4855 | 0.624 | 0.463 |
| | GATK | 0.128 | 0.535 | 0.207 | 0.002 | 0.004 | 0.002 | 0.065 | 0.2695 | 0.105 |
| | LYCEUM | 0.56 | 0.59 | 0.575 | 0.762 | 0.716 | 0.738 | 0.661 | 0.653 | 0.657 |
| 0.1x | CNVnator | 0.66 | 0.418 | 0.512 | 0.363 | 0.037 | 0.067 | 0.512 | 0.228 | 0.29 |
| | CONGA | 0.266 | 0.096 | 0.141 | 0.089 | 0.006 | 0.011 | 0.1775 | 0.051 | 0.076 |
| | Control-FREEC | 0.101 | 0.44 | 0.164 | 0.626 | 0.347 | 0.446 | 0.3635 | 0.3935 | 0.305 |
| | GATK | 0.069 | 0.423 | 0.119 | 0.0 | 0.001 | 0.0 | 0.0345 | 0.212 | 0.06 |
| | LYCEUM | 0.453 | 0.602 | 0.517 | 0.8 | 0.661 | 0.724 | 0.627 | 0.632 | 0.621 |
| 0.05x | CNVnator | 0.709 | 0.362 | 0.479 | 0.99 | 0.008 | 0.015 | 0.85 | 0.185 | 0.247 |
| | CONGA | 0.161 | 0.086 | 0.112 | 0.038 | 0.004 | 0.008 | 0.0995 | 0.045 | 0.06 |
| | Control-FREEC | 0.092 | 0.39 | 0.148 | 0.518 | 0.308 | 0.386 | 0.305 | 0.349 | 0.267 |
| | GATK | 0.052 | 0.408 | 0.093 | 0.0 | 0.001 | 0.0 | 0.026 | 0.2045 | 0.047 |
| | LYCEUM | 0.674 | 0.512 | 0.582 | 0.775 | 0.741 | 0.758 | 0.725 | 0.627 | 0.67 |

**Supplementary Table 2.** The performance comparison of the CNV callers on the down-sampled ancient samples (test set) in gene regions. CNVnator calls on original coverage samples are used as the ground truth.

| Coverage | Tools | DEL Precision | DEL Recall | DEL F1 Score | DUP Precision | DUP Recall | DUP F1 Score | Overall Precision | Overall Recall | Overall F1 Score |
|---|---|---|---|---|---|---|---|---|---|---|
| Original Coverage | CNVnator | 1.0 | 1.0 | 1.0 | 1.0 | 1.0 | 1.0 | 1.0 | 1.0 | 1.0 |
| | CONGA | 0.747 | 0.124 | 0.213 | 0.559 | 0.015 | 0.028 | 0.653 | 0.0695 | 0.1205 |
| | Control-FREEC | 0.2 | 0.741 | 0.315 | 0.867 | 0.786 | 0.824 | 0.534 | 0.764 | 0.57 |
| | GATK | 0.192 | 0.415 | 0.263 | 0.143 | 0.003 | 0.006 | 0.1675 | 0.209 | 0.1345 |
| | LYCEUM | 0.409 | 0.711 | 0.519 | 0.753 | 0.67 | 0.709 | 0.581 | 0.691 | 0.614 |
| 5x | CNVnator | 0.905 | 0.853 | 0.878 | 0.917 | 0.802 | 0.856 | 0.911 | 0.828 | 0.867 |
| | CONGA | 0.742 | 0.124 | 0.212 | 0.562 | 0.014 | 0.027 | 0.652 | 0.069 | 0.1195 |
| | Control-FREEC | 0.199 | 0.742 | 0.314 | 0.887 | 0.762 | 0.82 | 0.543 | 0.752 | 0.567 |
| | GATK | 0.272 | 0.339 | 0.302 | 0.054 | 0.002 | 0.004 | 0.163 | 0.1705 | 0.153 |
| | LYCEUM | 0.41 | 0.682 | 0.512 | 0.706 | 0.651 | 0.678 | 0.558 | 0.667 | 0.595 |
| 1x | CNVnator | 0.82 | 0.676 | 0.741 | 0.883 | 0.369 | 0.52 | 0.852 | 0.523 | 0.631 |
| | CONGA | 0.592 | 0.126 | 0.208 | 0.528 | 0.015 | 0.028 | 0.56 | 0.0705 | 0.118 |
| | Control-FREEC | 0.174 | 0.636 | 0.273 | 0.852 | 0.638 | 0.73 | 0.513 | 0.637 | 0.502 |
| | GATK | 0.267 | 0.317 | 0.29 | 0.0 | 0.0 | nan | 0.1335 | 0.1585 | 0.29 |
| | LYCEUM | 0.392 | 0.692 | 0.501 | 0.609 | 0.657 | 0.633 | 0.501 | 0.675 | 0.567 |
| 0.5x | CNVnator | 0.713 | 0.543 | 0.616 | 0.713 | 0.219 | 0.335 | 0.713 | 0.381 | 0.476 |
| | CONGA | 0.416 | 0.123 | 0.189 | 0.45 | 0.014 | 0.027 | 0.433 | 0.0685 | 0.108 |
| | Control-FREEC | 0.149 | 0.527 | 0.233 | 0.824 | 0.491 | 0.616 | 0.487 | 0.509 | 0.425 |
| | GATK | 0.231 | 0.314 | 0.266 | 0.004 | 0.004 | 0.004 | 0.1175 | 0.159 | 0.135 |
| | LYCEUM | 0.476 | 0.637 | 0.545 | 0.67 | 0.704 | 0.687 | 0.573 | 0.671 | 0.616 |
| 0.1x | CNVnator | 0.649 | 0.297 | 0.408 | 0.41 | 0.051 | 0.09 | 0.53 | 0.174 | 0.249 |
| | CONGA | 0.133 | 0.126 | 0.129 | 0.071 | 0.011 | 0.02 | 0.102 | 0.0685 | 0.0745 |
| | Control-FREEC | 0.099 | 0.335 | 0.153 | 0.55 | 0.276 | 0.367 | 0.325 | 0.306 | 0.26 |
| | GATK | 0.116 | 0.232 | 0.155 | 0.0 | 0.001 | 0.001 | 0.058 | 0.1165 | 0.078 |
| | LYCEUM | 0.422 | 0.636 | 0.507 | 0.742 | 0.656 | 0.696 | 0.582 | 0.646 | 0.602 |
| 0.05x | CNVnator | 0.68 | 0.259 | 0.375 | 1.0 | 0.013 | 0.026 | 0.84 | 0.136 | 0.201 |
| | CONGA | 0.078 | 0.109 | 0.091 | 0.03 | 0.008 | 0.012 | 0.054 | 0.0585 | 0.0515 |
| | Control-FREEC | 0.085 | 0.278 | 0.13 | 0.454 | 0.266 | 0.336 | 0.27 | 0.272 | 0.233 |
| | GATK | 0.081 | 0.231 | 0.12 | 0.001 | 0.001 | 0.001 | 0.041 | 0.116 | 0.0605 |
| | LYCEUM | 0.597 | 0.553 | 0.574 | 0.725 | 0.733 | 0.729 | 0.661 | 0.643 | 0.652 |

**Supplementary Table 3.** Confusion Matrices for the performance comparison of CNVnator on down-sampled ancient samples (test set) in exon regions. CNVnator calls on original coverage samples are used as the ground truth. These yield the precision and recall results for CNVnator in Supplementary Table 1.

| | | | Ground Truth | | |
|---|---|---|---|---|---|
| TOOL | Coverage | **Predicted** | NO CALL | DUP | DEL |
| CNVnator | Original Coverage | NO CALL | 1328288 | 0 | 0 |
| | | DUP | 0 | 12979 | 0 |
| | | DEL | 0 | 0 | 7542 |
| | 5x | NO CALL | 1326757 | 2253 | 657 |
| | | DUP | 1005 | 10717 | 0 |
| | | DEL | 526 | 9 | 6885 |
| | 1x | NO CALL | 1326400 | 6927 | 1665 |
| | | DUP | 827 | 6017 | 1 |
| | | DEL | 1061 | 35 | 5876 |
| | 0.5x | NO CALL | 1325884 | 9700 | 2482 |
| | | DUP | 1028 | 3266 | 2 |
| | | DEL | 1376 | 13 | 5058 |
| | 0.1x | NO CALL | 1325883 | 12448 | 4389 |
| | | DUP | 838 | 477 | 0 |
| | | DEL | 1567 | 54 | 3153 |
| | 0.05x | NO CALL | 1327186 | 12861 | 4813 |
| | | DUP | 1 | 99 | 0 |
| | | DEL | 1101 | 19 | 2729 |

**Supplementary Table 4.** Confusion Matrices for the performance comparison of CNVnator on down-sampled ancient samples (test set) in gene regions. CNVnator calls on original coverage samples are used as the ground truth. These yield the precision and recall results for CNVnator in Supplementary Table 2.

| | | | **Ground Truth** | | |
|---|---|---|---|---|---|
| TOOL | Coverage | **Predicted** | NO CALL | DUP | DEL |
| CNVnator | Original Coverage | NO CALL | 127963 | 0 | 0 |
| | | DUP | 0 | 1305 | 0 |
| | | DEL | 0 | 0 | 1093 |
| | 5x | NO CALL | 127786 | 242 | 161 |
| | | DUP | 95 | 1047 | 0 |
| | | DEL | 82 | 16 | 932 |
| | 1x | NO CALL | 127751 | 811 | 353 |
| | | DUP | 63 | 481 | 1 |
| | | DEL | 149 | 13 | 739 |
| | 0.5x | NO CALL | 127615 | 1014 | 499 |
| | | DUP | 114 | 286 | 1 |
| | | DEL | 234 | 5 | 593 |
| | 0.1x | NO CALL | 127705 | 1226 | 768 |
| | | DUP | 95 | 66 | 0 |
| | | DEL | 163 | 13 | 325 |
| | 0.05x | NO CALL | 127834 | 1284 | 810 |
| | | DUP | 0 | 17 | 0 |
| | | DEL | 129 | 4 | 283 |

**Supplementary Table 5.** Confusion Matrices for the performance comparison of CONGA on down-sampled ancient samples (test set) in exon regions. CNVnator calls on original coverage samples are used as the ground truth. These yield the precision and recall results for CONGA in Supplementary Table 1.

| | | | **Ground Truth** | | |
|---|---|---|---|---|---|
| TOOL | Coverage | **Predicted** | NO CALL | DUP | DEL |
| CONGA | Original Coverage | NO CALL | 1328062 | 12852 | 6842 |
| | | DUP | 103 | 115 | 0 |
| | | DEL | 123 | 12 | 700 |
| | 5x | NO CALL | 1328068 | 12862 | 6835 |
| | | DUP | 92 | 105 | 0 |
| | | DEL | 128 | 12 | 707 |
| | 1x | NO CALL | 1327966 | 12852 | 6808 |
| | | DUP | 113 | 115 | 0 |
| | | DEL | 209 | 12 | 734 |
| | 0.5x | NO CALL | 1327773 | 12862 | 6811 |
| | | DUP | 114 | 105 | 0 |
| | | DEL | 401 | 12 | 731 |
| | 0.1x | NO CALL | 1325498 | 12886 | 6816 |
| | | DUP | 802 | 78 | 0 |
| | | DEL | 1988 | 15 | 726 |
| | 0.05x | NO CALL | 1323509 | 12906 | 6896 |
| | | DUP | 1429 | 57 | 0 |
| | | DEL | 3350 | 16 | 646 |

**Supplementary Table 6.** Confusion Matrices for the performance comparison of CONGA on down-sampled ancient samples (test set) in gene regions. CNVnator calls on original coverage samples are used as the ground truth. These yield the precision and recall results for CONGA in Supplementary Table 2.

| | | | **Ground Truth** | | |
|---|---|---|---|---|---|
| TOOL | Coverage | **Predicted** | NO CALL | DUP | DEL |
| CONGA | Original Coverage | NO CALL | 127912 | 1276 | 957 |
| | | DUP | 15 | 19 | 0 |
| | | DEL | 36 | 10 | 136 |
| | 5x | NO CALL | 127912 | 1277 | 958 |
| | | DUP | 14 | 18 | 0 |
| | | DEL | 37 | 10 | 135 |
| | 1x | NO CALL | 127861 | 1276 | 955 |
| | | DUP | 17 | 19 | 0 |
| | | DEL | 85 | 10 | 138 |
| | 0.5x | NO CALL | 127763 | 1277 | 959 |
| | | DUP | 22 | 18 | 0 |
| | | DEL | 178 | 10 | 134 |
| | 0.1x | NO CALL | 126875 | 1280 | 955 |
| | | DUP | 195 | 15 | 0 |
| | | DEL | 893 | 10 | 138 |
| | 0.05x | NO CALL | 126242 | 1282 | 974 |
| | | DUP | 324 | 10 | 0 |
| | | DEL | 1397 | 13 | 119 |

**Supplementary Table 7.** Confusion Matrices for the performance comparison of Control-FREEC on down-sampled ancient samples (test set) in exon regions. CNVnator calls on original coverage samples are used as the ground truth. These yield the precision and recall results for Control-FREEC in Supplementary Table 1.

| | | | **Ground Truth** | | |
|---|---|---|---|---|---|
| TOOL | Coverage | **Predicted** | NO CALL | DUP | DEL |
| Control-FREEC | Original Coverage | NO CALL | 1297701 | 2483 | 1315 |
| | | DUP | 1771 | 10989 | 0 |
| | | DEL | 29212 | 29 | 6261 |
| | 5x | NO CALL | 1297972 | 2495 | 1293 |
| | | DUP | 1351 | 10858 | 1 |
| | | DEL | 29361 | 148 | 6282 |
| | 1x | NO CALL | 1297338 | 3911 | 1778 |
| | | DUP | 1716 | 9363 | 1 |
| | | DEL | 29630 | 227 | 5797 |
| | 0.5x | NO CALL | 1297449 | 5324 | 2607 |
| | | DUP | 1661 | 8010 | 4 |
| | | DEL | 29574 | 167 | 4965 |
| | 0.1x | NO CALL | 1296735 | 8279 | 4227 |
| | | DUP | 2785 | 4679 | 12 |
| | | DEL | 29164 | 543 | 3337 |
| | 0.05x | NO CALL | 1295987 | 8894 | 4597 |
| | | DUP | 3845 | 4159 | 27 |
| | | DEL | 28852 | 448 | 2952 |

**Supplementary Table 8.** Confusion Matrices for the performance comparison of Control-FREEC on down-sampled ancient samples (test set) in gene regions. CNVnator calls on original coverage samples are used as the ground truth. These yield the precision and recall results for Control-FREEC in Supplementary Table 2.

| | | | **Ground Truth** | | |
|---|---|---|---|---|---|
| TOOL | Coverage | **Predicted** | NO CALL | DUP | DEL |
| Control-FREEC | Original Coverage | NO CALL | 124588 | 257 | 282 |
| | | DUP | 155 | 1015 | 1 |
| | | DEL | 3213 | 20 | 809 |
| | 5x | NO CALL | 124598 | 279 | 282 |
| | | DUP | 125 | 985 | 0 |
| | | DEL | 3233 | 28 | 810 |
| | 1x | NO CALL | 124549 | 432 | 396 |
| | | DUP | 142 | 824 | 1 |
| | | DEL | 3265 | 36 | 695 |
| | 0.5x | NO CALL | 124566 | 630 | 515 |
| | | DUP | 135 | 635 | 1 |
| | | DEL | 3255 | 27 | 576 |
| | 0.1x | NO CALL | 124442 | 846 | 724 |
| | | DUP | 289 | 356 | 2 |
| | | DEL | 3225 | 90 | 366 |
| | 0.05x | NO CALL | 124356 | 864 | 784 |
| | | DUP | 409 | 344 | 4 |
| | | DEL | 3191 | 84 | 304 |

**Supplementary Table 9.** Confusion Matrices for the performance comparison of GATK on down-sampled ancient samples (test set) in exon regions. CNVnator calls on original coverage samples are used as the ground truth. These yield the precision and recall results for GATK in Supplementary Table 1.

| | | | **Ground Truth** | | |
|---|---|---|---|---|---|
| TOOL | Coverage | **Predicted** | NO CALL | DUP | DEL |
| GATK | Original Coverage | NO CALL | 1293972 | 2672 | 750 |
| | | DUP | 487 | 168 | 66 |
| | | DEL | 33411 | 9739 | 6620 |
| | 5x | NO CALL | 1309054 | 5979 | 2997 |
| | | DUP | 553 | 62 | 64 |
| | | DEL | 18263 | 6538 | 4375 |
| | 1x | NO CALL | 1306365 | 6539 | 3320 |
| | | DUP | 3234 | 28 | 0 |
| | | DEL | 18271 | 6012 | 4116 |
| | 0.5x | NO CALL | 1279198 | 6930 | 3454 |
| | | DUP | 27275 | 45 | 1 |
| | | DEL | 21397 | 5604 | 3981 |
| | 0.1x | NO CALL | 1241651 | 9291 | 4277 |
| | | DUP | 47275 | 12 | 12 |
| | | DEL | 38944 | 3276 | 3147 |
| | 0.05x | NO CALL | 1235763 | 10732 | 4398 |
| | | DUP | 39057 | 12 | 5 |
| | | DEL | 53050 | 1835 | 3033 |

**Supplementary Table 10.** Confusion Matrices for the performance comparison of GATK on down-sampled ancient samples (test set) in gene regions. CNVnator calls on original coverage samples are used as the ground truth. These yield the precision and recall results for GATK in Supplementary Table 2.

| | | | **Ground Truth** | | |
|---|---|---|---|---|---|
| TOOL | Coverage | **Predicted** | NO CALL | DUP | DEL |
| GATK | Original Coverage | NO CALL | 126471 | 857 | 639 |
| | | DUP | 24 | 4 | 0 |
| | | DEL | 1468 | 444 | 454 |
| | 5x | NO CALL | 127251 | 967 | 722 |
| | | DUP | 53 | 3 | 0 |
| | | DEL | 659 | 335 | 371 |
| | 1x | NO CALL | 127154 | 1018 | 747 |
| | | DUP | 147 | 0 | 0 |
| | | DEL | 662 | 287 | 346 |
| | 0.5x | NO CALL | 125912 | 1073 | 744 |
| | | DUP | 1137 | 5 | 6 |
| | | DEL | 914 | 227 | 343 |
| | 0.1x | NO CALL | 123505 | 1170 | 835 |
| | | DUP | 2655 | 1 | 4 |
| | | DEL | 1803 | 134 | 254 |
| | 0.05x | NO CALL | 123212 | 1225 | 835 |
| | | DUP | 1975 | 1 | 5 |
| | | DEL | 2776 | 79 | 253 |

**Supplementary Table 11.** Confusion Matrices for the performance comparison of LYCEUM on down-sampled ancient samples (test set) in exon regions. CNVnator calls on original coverage samples are used as the ground truth. These yield the precision and recall results for LYCEUM in Supplementary Table 1.

| | | | **Ground Truth** | | |
|---|---|---|---|---|---|
| TOOL | Coverage | **Predicted** | NO CALL | DUP | DEL |
| LYCEUM | Original Coverage | NO CALL | 1321674 | 3138 | 1784 |
| | | DUP | 1806 | 8620 | 232 |
| | | DEL | 4808 | 1221 | 5526 |
| | 5x | NO CALL | 1321129 | 2996 | 2155 |
| | | DUP | 2324 | 8548 | 259 |
| | | DEL | 4835 | 1435 | 5128 |
| | 1x | NO CALL | 1319088 | 3361 | 2369 |
| | | DUP | 2374 | 8097 | 260 |
| | | DEL | 6826 | 1521 | 4913 |
| | 0.5x | NO CALL | 1322792 | 3100 | 2772 |
| | | DUP | 2583 | 9298 | 321 |
| | | DEL | 2913 | 581 | 4449 |
| | 0.1x | NO CALL | 1321998 | 3280 | 2796 |
| | | DUP | 1941 | 8578 | 207 |
| | | DEL | 4349 | 1121 | 4539 |
| | 0.05x | NO CALL | 1323986 | 3092 | 3590 |
| | | DUP | 2704 | 9620 | 89 |
| | | DEL | 1598 | 267 | 3863 |

**Supplementary Table 12.** Confusion Matrices for the performance comparison of LYCEUM on down-sampled ancient samples (test set) in gene regions. CNVnator calls on original coverage samples are used as the ground truth. These yield the precision and recall results for LYCEUM in Supplementary Table 2.

| | | | **Ground Truth** | | |
|---|---|---|---|---|---|
| TOOL | Coverage | **Predicted** | NO CALL | DUP | DEL |
| LYCEUM | Original Coverage | NO CALL | 126764 | 256 | 280 |
| | | DUP | 251 | 875 | 36 |
| | | DEL | 948 | 174 | 777 |
| | 5x | NO CALL | 126767 | 264 | 311 |
| | | DUP | 317 | 850 | 37 |
| | | DEL | 879 | 191 | 745 |
| | 1x | NO CALL | 126463 | 267 | 296 |
| | | DUP | 509 | 858 | 41 |
| | | DEL | 991 | 180 | 756 |
| | 0.5x | NO CALL | 126898 | 280 | 350 |
| | | DUP | 406 | 919 | 47 |
| | | DEL | 659 | 106 | 696 |
| | 0.1x | NO CALL | 126906 | 282 | 373 |
| | | DUP | 273 | 856 | 25 |
| | | DEL | 784 | 167 | 695 |
| | 0.05x | NO CALL | 127269 | 287 | 474 |
| | | DUP | 348 | 957 | 15 |
| | | DEL | 346 | 61 | 604 |

**Supplementary Table 13.** The performance comparison of the CNV callers on simulated ancient samples (test set) in exon regions.

| **Coverage** | **Tools** | **DEL Precision** | **DEL Recall** | **DEL F1 Score** | **DUP Precision** | **DUP Recall** | **DUP F1 Score** | **Overall Precision** | **Overall Recall** | **Overall F1 Score** |
|---|---|---|---|---|---|---|---|---|---|---|
| 1x | CNVnator | 0.854 | 0.545 | 0.665 | 0.834 | 0.145 | 0.248 | 0.844 | 0.345 | 0.457 |
| | CONGA | 0.797 | 0.114 | 0.199 | 0.742 | 0.122 | 0.21 | 0.77 | 0.118 | 0.205 |
| | Control-FREEC | 0.027 | 0.144 | 0.045 | 0.852 | 0.03 | 0.058 | 0.44 | 0.087 | 0.052 |
| | LYCEUM | 0.924 | 0.811 | 0.864 | 0.942 | 0.949 | 0.945 | 0.933 | 0.88 | 0.905 |
| 0.5x | CNVnator | 0.79 | 0.381 | 0.514 | 0.666 | 0.029 | 0.055 | 0.728 | 0.205 | 0.285 |
| | CONGA | 0.747 | 0.111 | 0.194 | 0.738 | 0.12 | 0.206 | 0.743 | 0.116 | 0.2 |
| | Control-FREEC | 0.023 | 0.126 | 0.04 | 0.757 | 0.029 | 0.055 | 0.39 | 0.078 | 0.048 |
| | LYCEUM | 0.952 | 0.763 | 0.847 | 0.93 | 0.942 | 0.936 | 0.941 | 0.853 | 0.892 |
| 0.1x | CNVnator | 0.387 | 0.052 | 0.091 | 0.201 | 0.018 | 0.034 | 0.294 | 0.035 | 0.063 |
| | CONGA | 0.419 | 0.108 | 0.171 | 0.641 | 0.113 | 0.192 | 0.53 | 0.111 | 0.182 |
| | Control-FREEC | 0.022 | 0.12 | 0.038 | 0.558 | 0.023 | 0.045 | 0.29 | 0.072 | 0.042 |
| | LYCEUM | 0.922 | 0.778 | 0.844 | 0.934 | 0.929 | 0.932 | 0.928 | 0.854 | 0.888 |
| 0.05x | CNVnator | 0.616 | 0.025 | 0.049 | 0.175 | 0.004 | 0.007 | 0.396 | 0.015 | 0.028 |
| | CONGA | 0.275 | 0.102 | 0.149 | 0.556 | 0.109 | 0.182 | 0.416 | 0.106 | 0.166 |
| | Control-FREEC | 0.017 | 0.091 | 0.029 | 0.432 | 0.017 | 0.032 | 0.225 | 0.054 | 0.031 |
| | LYCEUM | 0.935 | 0.721 | 0.814 | 0.92 | 0.901 | 0.91 | 0.928 | 0.811 | 0.862 |

**Supplementary Table 14.** The performance comparison of the CNV callers on simulated ancient samples (test set) in gene regions.

| Coverage | Tools | DEL Precision | DEL Recall | DEL F1 Score | DUP Precision | DUP Recall | DUP F1 Score | Overall Precision | Overall Recall | Overall F1 Score |
|---|---|---|---|---|---|---|---|---|---|---|
| 1x | CNVnator | 0.839 | 0.438 | 0.576 | 0.916 | 0.096 | 0.173 | 0.878 | 0.267 | 0.375 |
| | CONGA | 0.753 | 0.174 | 0.283 | 0.86 | 0.142 | 0.244 | 0.807 | 0.158 | 0.264 |
| | Control-FREEC | 0.5 | 0.0 | 0.001 | 0.833 | 0.006 | 0.012 | 0.667 | 0.003 | 0.007 |
| | LYCEUM | 0.922 | 0.758 | 0.832 | 0.952 | 0.93 | 0.941 | 0.937 | 0.844 | 0.887 |
| 0.5x | CNVnator | 0.736 | 0.269 | 0.394 | 0.861 | 0.027 | 0.051 | 0.799 | 0.148 | 0.223 |
| | CONGA | 0.678 | 0.173 | 0.275 | 0.858 | 0.14 | 0.241 | 0.768 | 0.157 | 0.258 |
| | Control-FREEC | 0.5 | 0.0 | 0.001 | 0.442 | 0.007 | 0.014 | 0.471 | 0.004 | 0.008 |
| | LYCEUM | 0.952 | 0.712 | 0.815 | 0.927 | 0.921 | 0.924 | 0.94 | 0.817 | 0.87 |
| 0.1x | CNVnator | 0.207 | 0.01 | 0.02 | 0.48 | 0.024 | 0.046 | 0.344 | 0.017 | 0.033 |
| | CONGA | 0.315 | 0.17 | 0.221 | 0.711 | 0.133 | 0.225 | 0.513 | 0.152 | 0.223 |
| | Control-FREEC | 0.0 | 0.0 | 0.0 | 0.339 | 0.004 | 0.008 | 0.17 | 0.002 | 0.004 |
| | LYCEUM | 0.917 | 0.714 | 0.803 | 0.931 | 0.898 | 0.914 | 0.924 | 0.806 | 0.859 |
| 0.05x | CNVnator | 0.0 | 0.0 | 0.0 | 0.472 | 0.005 | 0.01 | 0.236 | 0.003 | 0.005 |
| | CONGA | 0.215 | 0.163 | 0.185 | 0.579 | 0.126 | 0.207 | 0.397 | 0.145 | 0.196 |
| | Control-FREEC | 0.0 | 0.0 | 0.0 | 0.571 | 0.003 | 0.006 | 0.286 | 0.002 | 0.003 |
| | LYCEUM | 0.923 | 0.682 | 0.785 | 0.905 | 0.869 | 0.886 | 0.914 | 0.776 | 0.836 |

**Supplementary Table 15.** Confusion Matrices for the performance comparison of CNVnator on simulated ancient samples (test set) in exon regions. These yield the precision and recall results for CNVnator in Supplementary Table 13.

| | | | Ground Truth | | |
|---|---|---|---|---|---|
| TOOL | Coverage | **Predicted** | NO CALL | DUP | DEL |
| CNVnator | 1x | NO CALL | 1881605 | 24403 | 6554 |
| | | DUP | 825 | 4150 | 0 |
| | | DEL | 1335 | 0 | 7838 |
| | 0.5x | NO CALL | 1881976 | 27652 | 8907 |
| | | DUP | 409 | 821 | 2 |
| | | DEL | 1380 | 80 | 5483 |
| | 0.1x | NO CALL | 1880539 | 28021 | 13630 |
| | | DUP | 2057 | 522 | 18 |
| | | DEL | 1169 | 10 | 744 |
| | 0.05x | NO CALL | 1883037 | 28446 | 14024 |
| | | DUP | 501 | 107 | 4 |
| | | DEL | 227 | 0 | 364 |

**Supplementary Table 16.** Confusion Matrices for the performance comparison of CNVnator on simulated ancient samples (test set) in gene regions. These yield the precision and recall results for CNVnator in Supplementary Table 14.

| | | | Ground Truth | | |
|---|---|---|---|---|---|
| TOOL | Coverage | **Predicted** | NO CALL | DUP | DEL |
| CNVnator | 1x | NO CALL | 178083 | 4410 | 1670 |
| | | DUP | 41 | 469 | 2 |
| | | DEL | 232 | 19 | 1304 |
| | 0.5x | NO CALL | 178066 | 4749 | 2175 |
| | | DUP | 21 | 130 | 0 |
| | | DEL | 269 | 19 | 801 |
| | 0.1x | NO CALL | 178114 | 4775 | 2945 |
| | | DUP | 128 | 118 | 0 |
| | | DEL | 114 | 5 | 31 |
| | 0.05x | NO CALL | 178318 | 4873 | 2976 |
| | | DUP | 28 | 25 | 0 |
| | | DEL | 10 | 0 | 0 |

**Supplementary Table 17.** Confusion Matrices for the performance comparison of CONGA on simulated ancient samples (test set) in exon regions. These yield the precision and recall results for CONGA in Supplementary Table 13.

| | | | **Ground Truth** | | |
|---|---|---|---|---|---|
| TOOL | Coverage | **Predicted** | NO CALL | DUP | DEL |
| CONGA | 1x | NO CALL | 1882136 | 25063 | 12752 |
| | | DUP | 1212 | 3490 | 0 |
| | | DEL | 417 | 0 | 1640 |
| | 0.5x | NO CALL | 1882012 | 25129 | 12787 |
| | | DUP | 1211 | 3424 | 2 |
| | | DEL | 542 | 0 | 1603 |
| | 0.1x | NO CALL | 1879816 | 25331 | 12842 |
| | | DUP | 1802 | 3221 | 2 |
| | | DEL | 2147 | 1 | 1548 |
| | 0.05x | NO CALL | 1877427 | 25438 | 12919 |
| | | DUP | 2480 | 3106 | 4 |
| | | DEL | 3858 | 9 | 1469 |

**Supplementary Table 18.** Confusion Matrices for the performance comparison of CONGA on simulated ancient samples (test set) in gene regions. These yield the precision and recall results for CONGA in Supplementary Table 14.

| | | | **Ground Truth** | | |
|---|---|---|---|---|---|
| TOOL | Coverage | **Predicted** | NO CALL | DUP | DEL |
| CONGA | 1x | NO CALL | 178121 | 4184 | 2426 |
| | | DUP | 82 | 697 | 31 |
| | | DEL | 153 | 17 | 519 |
| | 0.5x | NO CALL | 178045 | 4197 | 2432 |
| | | DUP | 83 | 685 | 30 |
| | | DEL | 228 | 16 | 514 |
| | 0.1x | NO CALL | 177047 | 4210 | 2445 |
| | | DUP | 242 | 653 | 24 |
| | | DEL | 1067 | 35 | 507 |
| | 0.05x | NO CALL | 176194 | 4243 | 2466 |
| | | DUP | 425 | 618 | 25 |
| | | DEL | 1737 | 37 | 485 |

**Supplementary Table 19.** Confusion Matrices for the performance comparison of Control-FREEC on simulated ancient samples (test set) in exon regions. These yield the precision and recall results for Control-FREEC in Supplementary Table 13.

| | | | **Ground Truth** | | |
|---|---|---|---|---|---|
| TOOL | Coverage | **Predicted** | NO CALL | DUP | DEL |
| Control-FREEC | 1x | NO CALL | 1809002 | 28061 | 12331 |
| | | DUP | 147 | 868 | 4 |
| | | DEL | 75386 | 0 | 2081 |
| | 0.5x | NO CALL | 1808909 | 28101 | 12604 |
| | | DUP | 264 | 826 | 1 |
| | | DEL | 75362 | 2 | 1811 |
| | 0.1x | NO CALL | 1808363 | 28246 | 12693 |
| | | DUP | 537 | 678 | 0 |
| | | DEL | 75635 | 5 | 1723 |
| | 0.05x | NO CALL | 1808029 | 28445 | 13099 |
| | | DUP | 635 | 482 | 0 |
| | | DEL | 75871 | 2 | 1317 |

**Supplementary Table 20.** Confusion Matrices for the performance comparison of Control-FREEC on simulated ancient samples (test set) in gene regions. These yield the precision and recall results for Control-FREEC in Supplementary Table 14.

| | | | Ground Truth | | |
|---|---|---|---|---|---|
| TOOL | Coverage | **Predicted** | NO CALL | DUP | DEL |
| Control-FREEC | 1x | NO CALL | 178319 | 4868 | 2975 |
| | | DUP | 6 | 30 | 0 |
| | | DEL | 1 | 0 | 1 |
| | 0.5x | NO CALL | 178282 | 4864 | 2975 |
| | | DUP | 43 | 34 | 0 |
| | | DEL | 1 | 0 | 1 |
| | 0.1x | NO CALL | 178289 | 4879 | 2976 |
| | | DUP | 37 | 19 | 0 |
| | | DEL | 0 | 0 | 0 |
| | 0.05x | NO CALL | 178314 | 4882 | 2976 |
| | | DUP | 12 | 16 | 0 |
| | | DEL | 0 | 0 | 0 |

**Supplementary Table 21.** Confusion Matrices for the performance comparison of LYCEUM on simulated ancient samples (test set) in exon regions. These yield the precision and recall results for LYCEUM in Supplementary Table 13.

| | | | Ground Truth | | |
|---|---|---|---|---|---|
| TOOL | Coverage | **Predicted** | NO CALL | DUP | DEL |
| LYCEUM | 1x | NO CALL | 1881256 | 1422 | 2625 |
| | | DUP | 1580 | 27096 | 88 |
| | | DEL | 929 | 35 | 11679 |
| | 0.5x | NO CALL | 1881315 | 1659 | 3285 |
| | | DUP | 1902 | 26883 | 120 |
| | | DEL | 548 | 11 | 10987 |
| | 0.1x | NO CALL | 1881113 | 1970 | 3106 |
| | | DUP | 1767 | 26521 | 94 |
| | | DEL | 885 | 62 | 11192 |
| | 0.05x | NO CALL | 1880964 | 2763 | 3912 |
| | | DUP | 2144 | 25728 | 103 |
| | | DEL | 657 | 62 | 10377 |

**Supplementary Table 22.** Confusion Matrices for the performance comparison of LYCEUM on simulated ancient samples (test set) in gene regions. These yield the precision and recall results for LYCEUM in Supplementary Table 14.

| | | | Ground Truth | | |
|---|---|---|---|---|---|
| TOOL | Coverage | **Predicted** | NO CALL | DUP | DEL |
| LYCEUM | 1x | NO CALL | 177974 | 328 | 694 |
| | | DUP | 205 | 4556 | 26 |
| | | DEL | 177 | 14 | 2256 |
| | 0.5x | NO CALL | 177950 | 380 | 810 |
| | | DUP | 307 | 4510 | 47 |
| | | DEL | 99 | 8 | 2119 |
| | 0.1x | NO CALL | 177888 | 476 | 823 |
| | | DUP | 298 | 4400 | 28 |
| | | DEL | 170 | 22 | 2125 |
| | 0.05x | NO CALL | 177797 | 627 | 904 |
| | | DUP | 407 | 4254 | 41 |
| | | DEL | 152 | 17 | 2031 |

**Supplementary Table 23.** The performance comparison of LYCEUM on 7 moderate coverage real ancient samples samples at their original coverage, categorized by their contamination levels. The c values indicate contamination predictions obtained using ContaminationX. Refer to Supplementary Table 24 for the detailed ContaminationX output for the test set.

| | DEL Precision | DEL Recall | DEL F1 | DUP Precision | DUP Recall | DUP F1 | Overall Precision | Overall Recall | Overall F1 |
|---|---|---|---|---|---|---|---|---|---|
| Low Contamination ($c < 1\%$) | 0.398 | 0.788 | 0.529 | 0.825 | 0.664 | 0.736 | 0.611 | 0.726 | 0.632 |
| High Contamination ($c > 30\%$) | 0.594 | 0.686 | 0.637 | 0.788 | 0.665 | 0.721 | 0.691 | 0.675 | 0.679 |

**Supplementary Table 24.** ContaminationX output for 7 moderate coverage real ancient samples at their original coverage levels.

| Sample Name | Method | ContaminationEstimate | LowerCI | UpperCI | ErrorRate | SitesUsed |
|---|---|---|---|---|---|---|
| Nea3 | One-cns | 0.3924 | 0.3919 | 0.3930 | 0.0011 | 55756 |
| | Two-cns | 0.3974 | 0.3969 | 0.3979 | 0.0011 | 55756 |
| AKT16 | One-cns | 0.3697 | 0.3691 | 0.3703 | 0.0012 | 52695 |
| | Two-cns | 0.3779 | 0.3773 | 0.3784 | 0.0012 | 52695 |
| STAR1 | One-cns | 0.3665 | 0.3660 | 0.3670 | 0.0011 | 56199 |
| | Two-cns | 0.3736 | 0.3731 | 0.3741 | 0.0011 | 56199 |
| VLASA7 | One-cns | 0.0057 | 0.0056 | 0.0058 | 0.0009 | 55396 |
| | Two-cns | 0.0058 | 0.0057 | 0.0059 | 0.0009 | 55396 |
| Bichon | One-cns | 0.0031 | 0.0029 | 0.0032 | 0.0029 | 38725 |
| | Two-cns | 0.0032 | 0.0030 | 0.0033 | 0.0029 | 38725 |
| VLASA32 | One-cns | 0.0064 | 0.0063 | 0.0065 | 0.0012 | 51731 |
| | Two-cns | 0.0064 | 0.0064 | 0.0065 | 0.0012 | 51731 |
| Ust | One-cns | 0.0040 | 0.0040 | 0.0041 | 0.0013 | 37174 |
| | Two-cns | 0.0040 | 0.0040 | 0.0041 | 0.0013 | 37174 |

**Supplementary Table 25.** Metadata for 13 moderate-coverage ancient samples used in the fine-tuning of LYCEUM. [1–7].

| Sample_ID | ID_publication | Population Name | Country | Data Type | Publication | Genome Coverage | Date | Latitude | Longitude |
|---|---|---|---|---|---|---|---|---|---|
| **Anzick-1** | Anzick-1 | Anzick-1 | USA | Shotgun | RasmussenNature2014 | 11,915 | 10776-10622 calBCE | 45,993056 | -110,66139 |
| **BAR25** | BAR25 | Anatolia_N | Turkey | Shotgun | MarchibioRxiv2020 | 12,65 | 8384-8205 calBP | 40,3 | 29,56666 |
| **BOT2016** | BOT2016 | Botai | Kazakhstan | Shotgun | DamgaardScience2018 | 12,991 | 3632-3368 calBCE (4695 ±50 BP, UBA-32666) | 53,305983 | 67,648166 |
| **irk034** | irk034 | Siberia_Cis_Baikal_LN | Russia | Shotgun | KılınçSciAdv2021 | 14,5 | 3645-3521 calBCE | 53,2206944 | 103,39475 |
| **Kotias** | Kotias | CHG | Georgia | Shotgun | JonesNatureCommunications2015 | 11,5591 | 7940-7600 calBCE [7938-7580 calBCE (8665 ±65 BP, RTT-5246), 7946-7612 calBCE (8745 ±40, OxA-28256)] | 42,28 | 43,28 |
| **LBK** | LBK | Stuttgart.SG | Germany | Shotgun | LazaridisNature2014 | 14,267 | 5310-5070 calBCE (6246 ±30 BP, MAMS-24635) | 48,78 | 9,18 |
| **LEPE48** | LEPE48 | Iron_Gates_HG_EN | Serbia | Shotgun | MarchibioRxiv2020 | 10,92 | 8012-7867 calBP | 44,552924 | 22,027563 |
| **LEPE52** | LEPE52 | Iron Gates E-MN | Serbia | Shotgun | MarchibioRxiv2020 | 12,37 | 7931-7693 calBP | 44,552924 | 22,027563 |
| **Loschbour** | Loschbour_published.DG | Loschbour.SG | Luxembourg | Shotgun | LazaridisNature2014 | 15,526 | 6210-5990 calBCE (7205 ±50 BP, OxA-7738) | 49,81 | 6,4 |
| **Nea2** | Nea2 | Greece_N | Greece | Shotgun | MarchibioRxiv2020 | 12,51 | 8173-8023 calBP | 40,5879 | 22,2512 |
| **VC3-2** | VC3-2 | Starcevo_EN | Serbia | Shotgun | MarchibioRxiv2020 | 11,22 | 7565-7426 calBP | 44,762 | 20,6232 |
| **WC1** | WC1 | Iran_WezmehCave_N | Iran | Shotgun | BroushakiScience2016 | 9,15617 | 7455-7082 calBCE (8240 ±56 BP, UBA-25840) | 34,6129 | 47,1057 |
| **Yamnaya** | Yamnaya | Russia_EBA | Russia | Shotgun | DamgaardScience2018 | 23,312 | 3018-2887 calBCE (4315 ±34 BP, UBA-32667) | 49,134217 | 75,851817 |

**Supplementary Table 26.** Metadata for 7 moderate coverage ancient samples used in Section 2.4 to evaluate and compare the performance of LYCEUM with other methods [2, 5, 8].

| Sample_ID | ID_publication | Population Name | Country | Data Type | Publication | Genome Coverage | Date | Latitude | Longitude |
|---|---|---|---|---|---|---|---|---|---|
| **AKT16** | AKT16 | Anatolia_EN | Turkey | Shotgun | MarchibioRxiv2020 | 12,25 | 8635-8460 calBP | 40,16707 | 28,7756103 |
| **Bichon** | Bichon.SG | Bichon | Switzerland | Shotgun | JonesNatureCommunications2015 | 13,521 | 11820-11610 calBCE (11855 ±50 BP, OxA-27763) | 47,0999985 | 6,86999989 |
| **Nea3** | Nea3 | Greece_N | Greece | Shotgun | MarchibioRxiv2020 | 11,57 | 8327-8040 calBP | 40,5879 | 22,2512 |
| **STAR1** | STAR1 | Starcevo_EN | Serbia | Shotgun | MarchibioRxiv2020 | 10,55 | 7589-7476 calBP | 44,8218 | 20,7299 |
| **Ust** | Ust_Ishim_published.DG | Ust_Ishim | Russia | Shotgun | FuNature2014 | 26,345 | 45530-40610 calBCE [46064-40920 calBCE (41400 ±1300 BP, OxA-25516), 46364-40844 calBCE (41400 ±1400 BP, OxA-30190)] | 57,7 | 71,1 |
| **VLASA32** | VLASA32 | Iron_Gates_HG | Serbia | Shotgun | MarchibioRxiv2020 | 12,65 | 9741-9468 calBP | 44,53 | 22,05 |
| **VLASA7** | VLASA7 | Iron Gates LM | VLASA7 | Shotgun | MarchibioRxiv2020 | 15,21 | 8764-8340 calBP | 44,53 | 22,05 |

**Supplementary Table 27.** Metadata for 50 ancient samples used in analyses described in Section 2.5 and 2.6 [1–19]

| Sample_ID | ID_publication | Population Name | Country | Data Type | Publication | Genome Coverage | Date | Latitude | Longitude |
|---|---|---|---|---|---|---|---|---|---|
| **AKT16** | AKT16 | Anatolia_EN | Turkey | Shotgun | MarchibioRxiv2020 | 12,25 | 8635-8460 calBP | 40,16707 | 28,77561 |
| **Ash128** | Ash128 | Anatolia_EN | Turkey | Shotgun | YakaCurrentBiology2021 | 5,03217 | 8240-7941 (%95,4) | 38,34 | 34,23 |
| **ATP2** | ATP2.SG | Iberia_Chl | Spain | Shotgun | GuntherPNAS2015 | 4,080 | 2900-2679 calBCE (4210 ±30 BP, Beta-386394) | 42,3525 | -3,51833 |
| **Bar31** | Bar31.SG | Anatolian_N | Turkey | Shotgun | HofmanovaPNAS2016 | 3,374 | 6419-6238 calBCE (7457 ±44 BP, UBA-29838) | 40,3 | 29,56667 |
| **Bar8** | Bar8.SG | Anatolian_N | Turkey | Shotgun | HofmanovaPNAS2016 | 6,285 | 6212-6030 calBCE (7238 ±38 BP, UBA-29837) | 40,3 | 29,56667 |
| **Bichon** | Bichon.SG | Bichon | Switzerland | Shotgun | JonesNatureCommunications2015 | 13,521 | 11820-11610 calBCE (11855 ±50 BP, OxA-27763) | 47,1 | 6,87 |
| **Bon002** | Bon002.SG | Boncuklu | Turkey | Whole Genome Capture | KilincCurrentBiology2016 | 6,688 | 8279-7977 calBCE (Wk-29763) | 37,75191 | 32,8649 |
| **BOT14** | BOT14 | Botai | Kazakhstan | Shotgun | DamgaardScience2018 | 3,570 | 3518-3109 calBCE (4598 ±46 BP, UBA-32662) | 53,30598 | 67,64817 |
| **BOT15** | BOT15 | Botai | Kazakhstan | Shotgun | DamgaardScience2018 | 2,895 | 3343-3026 calBCE (4474 ±37 BP, UBA-32663) | 53,30598 | 67,64817 |
| **Chan** | Chan.SG | Iberia_HG | Spain | Shotgun | GonzalesFortesCurrentBiology2017 | 1,580 | 7305-7057 calBCE (8155 ±42 BP, Ua-13398/Ua-38115) | 43,14516 | -7,04093 |
| **CovaMoura9B** | CM9B | Portugal_LN_Chl | Portugal | Shotgun | MartinianoPLoSGenetics2017 | 2,25947 | 3700-2200 BCE | 38,74504 | -9,21522 |
| **DA100** | DA100 | TianShanHun | Kyrgyzstan | Shotgun | DamgaardNature2018 | 1,902 | 178-423 calCE (1719 ±48 BP, UBA-31213) | 42,15667 | 77,40556 |
| **DA222** | DA222 | Karluk | Kyrgyzstan | Shotgun | DamgaardNature2018 | 3,269 | 750-950 CE | 43,2025 | 76,98167 |
| **DA243** | DA243 | Alan | Russia | Shotgun | DamgaardNature2018 | 2,780 | 450-1350 CE | 43,95865 | 42,5873 |
| **DA249** | DA249 | Shamanka_EN | Russia | Shotgun | DamgaardScience2018 | 4,374 | 5988-5791 calBCE (7005 ±40 BP, OxA-21535) | 51,69833 | 103,7031 |
| **DA28** | DA28 | GoldenHordeAsian | Kazakhstan | Shotgun | DamgaardNature2018 | 3,682 | 1200-1600 CE | 46,99722 | 66,28583 |
| **DA379** | DA379 | Namazga_CA | Turkmenistan | Shotgun | DamgaardScience2018 | 0,0434962 | 3370-3023 calBCE (4515 ±57 BP, UBA-33658) | 37,60098 | 59,32841 |
| **DA380** | DA380 | Namazga_CA | Turkmenistan | Shotgun | DamgaardScience2018 | 0,444332 | 3364-3097 calBCE (4528 ±40 BP, UBA-33659) | 37,60098 | 59,32841 |
| **DA45** | DA45 | XiongNu | Mongolia | Shotgun | DamgaardNature2018 | 7,960 | 183-41 calBCE (2083 ±27 BP, UBA-31155) | 42,52583 | 105,18 |
| **EBA1** | EBA1 | Central_Steppe_EMBA | Russia | Shotgun | DamgaardScience2018 | 3,885 | 2286-2037 calBCE (3752 ±35 BP, UBA-32665) | 51,63134 | 74,66445 |
| **EBA2** | EBA2 | Central_Steppe_EMBA | Russia | Shotgun | DamgaardScience2018 | 8,814 | 2620-2468 calBCE (4013 ±33 BP, UBA-32664) | 52,62978 | 76,72765 |
| **GB** | GB.SG | Iron_Gates_EN | Romania | Shotgun | GonzalesFortesCurrentBiology2017 | 3,370 | 3506-3349 calBCE (4621 ±28 BP, MAMS-28614) | 46,52616 | 26,92981 |
| **Klei10** | Klei10 | Greece_Klei10 | Greece | Shotgun | HofmanovaPNAS2016 | 1,91646 | 4230-3995 calBCE (5559 ±22 BP, MAMS-23038) | 40,43 | 21,78 |
| **LugarCanto42** | LC42 | Portugal_MN | Portugal | Shotgun | MartinianoPLoSGenetics2017 | 2,60478 | 4500-3500 BCE | 39,00812 | 9,390754 |
| **LugarCanto44** | LC44 | Portugal_MN | Portugal | Shotgun | MartinianoPLoSGenetics2017 | 1,77033 | 4500-3500 BCE | 39,00812 | 9,390754 |
| **Motala12** | Motala12 | Motala12_SG | Sweden | Shotgun | LazaridisNature2014 | 1,92948 | 5722-5628 calBCE (6773 ±30 BP, Ua-51723) | 58,535 | 15,046 |
| **Nea3** | Nea3 | Greece_N | Greece | Shotgun | MarchibioRxiv2020 | 11,57 | 8327-8040 calBP | 40,5879 | 22,2512 |
| **NEO240** | NEO240 | Neolithic | Russia | Shotgun | SikoraNature2019 | 6,16589 | 5625-5481 calBCE (6627 ±45 BP, UBA-33768) | 44,5 | 135,4 |
| **R1** | R1 | Italian_LBA-EIA | Italy | Shotgun | AntonioScience2019 | 3,669 | 930 - 839 calBCE | 42,88014 | 13,89377 |
| **R2** | R2 | Italian_EN | Italy | Shotgun | AntonioScience2019 | 3,330 | 6066 - 6002 calBCE | 41,96 | 13,54 |
| **R3** | R3 | Italian_UPP | Italy | Shotgun | AntonioScience2019 | 3,702 | 5836 - 5723 calBCE | 41,96 | 13,54 |
| **R4** | R4 | Italian_UPN | Italy | Shotgun | AntonioScience2019 | 3,309 | 2950 - 2880 calBCE | 41,96 | 13,54 |
| **R7** | R7 | Italian_UPN | Italy | Shotgun | AntonioScience2019 | 2,673 | 8821 - 8642 calBCE | 41,96 | 13,54 |
| **R9** | R9 | Italian_EN | Italy | Shotgun | AntonioScience2019 | 3,453 | 5607 - 5485 calBCE | 41,96 | 13,54 |
| **RISE493** | RISE493 | Karasut | Russia | Shotgun | AllentoftNature2015 | 4,972 | 1531-1427 calBCE (3214 ±26 BP, OxA-31211) | 53,152 | 91,05 |
| **RISE495** | RISE495 | Karasut | Russia | Shotgun | AllentoftNature2015 | 2,996 | 1400-900 BCE | 52,954 | 90,187 |
| **RISE496** | RISE496 | Karasut | Russia | Shotgun | AllentoftNature2015 | 1,599 | 1414-1261 calBCE (3070 ±28 BP, OxA-31213) | 52,954 | 90,187 |
| **RISE497** | RISE497 | Karasut | Russia | Shotgun | AllentoftNature2015 | 5,507 | 1400-900 BCE | 52,954 | 90,187 |
| **RISE505** | RISE505 | Adronovo | Russia | Shotgun | AllentoftNature2015 | 3,482 | 1746-1626 calBCE (3391 ±27 BP, OxA-31216) | 53,456 | 85,447 |
| **RISE511** | RISE511 | Afanasievo | Russia | Shotgun | AllentoftNature2015 | 2,448 | 2909-2679 calBCE (4224 ±36 BP, OxA-31568) | 54,584 | 90,775 |
| **RISE664** | RISE664 | Okunevo_EMBA | Russia | Shotgun | DamgaardScience2018 | 4,261 | 2459-2206 calBCE (3850 ±38 BP, UBA-31593) | 53,54781 | 91,02565 |
| **RISE675** | RISE675 | Okunevo_EMBA | Russia | Shotgun | DamgaardScience2018 | 0,503469 | 2859-2350 calBCE (4023 ±56 BP, UBA-31597) | 53,70856 | 90,35981 |
| **RISE98** | RISE98 | Nordic_LN | Sweden | Shotgun | AllentoftNature2015 | 4,670 | 2275-2032 calBCE (3736 ±32 BP, OxA-28987) | 55,381 | 13,445 |
| **SC2** | SC2 | Iron_Gates_HG.SG | Serbia-romania | Shotgun | GonzalesFortesCurrentBiology2017 | 2,69884 | 7250-6500 BCE | 44,62971 | 22,61256 |
| **scy009** | scy009 | Scythian | Ukraine | Shotgun | KrzewińskaScienceAdvances2018 | 2,11486 | 768-431 calBCE (2470 ±30 BP, Beta-451570) | 46,33283 | 29,46166 |
| **STAR1** | STAR1 | Starcevo_EN | Serbia | Shotgun | MarchibioRxiv2020 | 10,55 | 7589-7476 calBP | 44,8218 | 20,7299 |
| **Ust** | Ust_Ishim_published.DG | Ust_Ishim | Russia | Shotgun | FuNature2014 | 26,345 | 45530-40610 calBCE [46064-40920 calBCE (41400 ±1300 BP, OxA-25516), 46364-40844 calBCE (41400 ±1400 BP, OxA-30190)] | 57,7 | 71,1 |
| **VLASA32** | VLASA32 | Iron_Gates_HG | Serbia | Shotgun | MarchibioRxiv2020 | 12,65 | 9741-9468 calBP | 44,53 | 22,05 |
| **VLASA7** | VLASA7 | Iron Gates LM | VLASA7 | Shotgun | MarchibioRxiv2020 | 15,21 | 8764-8340 calBP | 44,53 | 22,05 |
| **Yana** | Yana1 | Upper-Paleolithic | Russia | Shotgun | SikoraNature2019 | 20,4714 | 30250-29550 BCE | 70,7 | 135,4 |

**Supplementary Table 28.** List of 550 samples from the 1000 Genomes dataset selected for pre-training LYCEUM.

| | | | | | | | | | |
|---|---|---|---|---|---|---|---|---|---|
| HG00472 | HG01119 | HG02190 | HG01444 | HG01113 | HG00288 | HG01860 | HG00237 | HG02023 | HG02356 |
| HG02292 | HG01809 | HG00622 | HG02239 | HG01918 | HG01066 | HG01372 | HG01048 | HG01173 | HG01353 |
| HG01468 | HG00336 | HG02089 | HG01359 | HG01606 | HG01869 | HG00879 | HG00867 | HG01111 | HG01474 |
| HG02079 | HG01075 | HG01857 | HG02139 | HG02230 | HG01531 | HG00473 | HG02332 | HG01051 | HG01164 |
| HG01605 | HG01072 | HG01198 | HG01865 | HG01253 | HG01845 | HG00736 | HG01354 | HG01802 | HG01610 |
| HG01187 | HG01801 | HG01049 | HG01110 | HG01485 | HG00253 | HG01598 | HG01926 | HG01088 | HG01921 |
| HG01398 | HG02182 | HG01360 | HG02231 | HG00421 | HG00554 | HG01497 | HG01190 | HG00242 | HG00631 |
| HG01847 | HG01351 | HG01783 | HG01271 | HG02262 | HG01844 | HG00331 | HG00304 | HG02307 | HG00442 |
| HG01914 | HG01461 | HG00096 | HG01256 | HG01507 | HG01811 | HG01489 | HG02143 | HG01509 | HG00662 |
| HG01069 | HG01756 | HG01492 | HG02142 | HG00252 | HG00864 | HG01956 | HG00277 | HG00419 | HG01396 |
| HG00332 | HG01437 | HG00367 | HG01272 | HG02238 | HG00260 | HG02156 | HG00251 | HG00614 | HG02233 |
| HG02075 | HG01880 | HG01958 | HG01086 | HG01063 | HG01628 | HG01840 | HG01241 | HG00590 | HG00674 |
| HG02069 | HG00844 | HG00268 | HG01047 | HG01571 | HG01341 | HG01205 | HG00326 | HG00632 | HG01055 |
| HG00250 | HG02113 | HG01870 | HG01768 | HG02072 | HG02186 | HG01085 | HG01684 | HG00099 | HG01124 |
| HG00583 | HG01858 | HG01572 | HG01438 | HG00422 | HG01747 | HG02090 | HG02298 | HG00325 | HG02166 |
| HG01623 | HG00478 | HG00436 | HG00543 | HG01257 | HG01149 | HG01920 | HG01174 | HG01441 | HG00640 |
| HG00278 | HG02317 | HG01710 | HG02259 | HG02061 | HG00449 | HG02084 | HG02028 | HG02006 | HG01046 |
| HG01200 | HG01139 | HG02009 | HG01350 | HG02085 | HG01894 | HG00464 | HG01583 | HG02284 | HG02019 |
| HG02291 | HG02215 | HG01125 | HG01402 | HG01848 | HG01095 | HG02282 | HG01518 | HG01079 | HG02017 |
| HG00334 | HG00309 | HG01953 | HG01133 | HG01284 | HG01597 | HG01501 | HG00310 | HG02322 | HG02047 |
| HG01630 | HG01204 | HG00382 | HG00372 | HG00653 | HG01915 | HG01326 | HG00353 | HG01512 | HG01369 |
| HG02285 | HG00245 | HG00551 | HG00592 | HG00634 | HG00276 | HG01248 | HG00692 | HG00280 | HG02337 |
| HG01893 | HG00537 | HG02081 | HG01494 | HG01777 | HG02140 | HG02051 | HG01889 | HG01440 | HG00607 |
| HG01790 | HG01140 | HG01805 | HG01521 | HG00536 | HG01699 | HG01176 | HG01323 | HG01362 | HG01816 |
| HG01851 | HG01625 | HG01631 | HG00542 | HG02070 | HG00598 | HG00595 | HG00463 | HG02050 | HG00239 |
| HG01412 | HG02188 | HG02064 | HG02278 | HG00256 | HG02088 | HG01191 | HG00324 | HG01992 | HG01971 |
| HG00663 | HG00375 | HG00327 | HG01762 | HG00534 | HG01708 | HG01890 | HG01771 | HG00560 | HG02016 |
| HG02082 | HG01305 | HG01668 | HG00445 | HG00384 | HG02179 | HG02185 | HG01134 | HG01935 | HG02057 |
| HG00448 | HG01613 | HG00361 | HG01912 | HG00345 | HG00369 | HG01148 | HG01524 | HG01593 | HG01990 |
| HG02151 | HG01991 | HG01058 | HG01551 | HG01694 | HG01137 | HG00357 | HG01413 | HG02147 | HG01879 |
| HG01883 | HG00531 | HG01131 | HG01927 | HG01988 | HG00257 | HG02223 | HG02283 | HG01866 | HG02353 |
| HG01941 | HG01686 | HG02232 | HG00254 | HG01171 | HG01578 | HG00623 | HG00376 | HG01259 | HG01797 |
| HG00321 | HG01974 | HG00732 | HG00672 | HG02138 | HG01702 | HG02054 | HG00263 | HG00360 | HG01950 |
| HG02277 | HG01980 | HG00329 | HG01308 | HG02343 | HG01516 | HG02309 | HG01101 | HG01620 | HG00479 |
| HG01073 | HG01393 | HG00404 | HG00306 | HG02219 | HG02031 | HG01603 | HG02136 | HG02104 | HG01938 |
| HG01789 | HG01488 | HG02058 | HG00328 | HG01456 | HG01281 | HG00701 | HG00619 | HG00475 | HG00625 |
| HG00271 | HG00378 | HG01348 | HG00368 | HG01673 | HG00629 | HG01432 | HG01932 | HG00656 | HG02102 |
| HG01600 | HG01595 | HG01112 | HG00362 | HG00476 | HG00265 | HG01612 | HG00699 | HG01682 | HG01849 |
| HG02299 | HG01976 | HG01817 | HG00246 | HG00236 | HG01334 | HG02165 | HG00525 | HG01356 | HG01530 |
| HG01061 | HG01800 | HG01810 | HG01108 | HG01345 | HG00338 | HG01363 | HG00315 | HG00234 | HG00690 |
| HG02272 | HG01779 | HG00410 | HG02351 | HG00596 | HG00318 | HG00452 | HG01504 | HG02060 | HG01757 |
| HG00743 | HG01269 | HG01250 | HG01565 | HG01589 | HG01121 | HG00500 | HG01704 | HG00599 | HG02134 |
| HG00232 | HG02312 | HG01525 | HG02086 | HG01162 | HG01806 | HG01871 | HG02116 | HG01550 | HG01566 |
| HG01855 | HG00740 | HG01965 | HG00684 | HG01945 | HG00978 | HG02152 | HG01617 | HG01031 | HG01197 |
| HG01670 | HG01130 | HG01375 | HG01979 | HG01462 | HG01342 | HG02153 | HG00610 | HG01776 | HG00341 |
| HG01970 | HG01455 | HG00255 | HG02266 | HG00379 | HG01495 | HG01815 | HG01384 | HG02256 | HG00355 |
| HG01459 | HG00628 | HG00956 | HG01029 | HG01182 | HG02220 | HG01798 | HG01168 | HG02330 | HG00356 |
| HG02286 | HG00346 | HG01812 | HG02315 | HG01503 | HG01092 | HG00313 | HG02155 | HG01924 | HG00657 |
| HG01968 | HG01344 | HG00350 | HG01794 | HG01161 | HG00766 | HG00611 | HG02073 | HG01951 | HG02164 |
| HG01142 | HG00409 | HG00403 | HG00698 | HG02053 | HG00238 | HG00581 | HG01107 | HG02187 | HG01765 |
| HG01859 | HG01882 | HG01868 | HG00351 | HG01506 | HG02128 | HG01537 | HG02040 | HG01357 | HG00281 |
| HG02250 | HG01302 | HG00428 | HG00593 | HG00290 | HG02137 | HG02325 | HG01105 | HG01077 | HG00734 |
| HG01491 | HG02314 | HG02002 | HG00566 | HG01247 | HG01709 | HG02095 | HG02048 | HG00689 | HG02265 |
| HG00330 | HG00358 | HG02076 | HG01447 | HG00097 | HG01556 | HG00261 | HG01136 | HG00373 | HG01365 |
| HG01094 | HG01863 | HG01104 | HG01700 | HG01936 | HG02345 | HG01366 | HG00443 | HG00337 | HG01389 |
